# Dissecting Conformational Changes in APP’s Transmembrane Domain Linked to ε-Efficiency in Familial Alzheimer’s Disease

**DOI:** 10.1101/269084

**Authors:** Alexander Götz, Christina Scharnagl

## Abstract

The mechanism by which familial Alzheimer’s disease (FAD) mutations within the transmembrane domain (TMD) of the Amyloid Precursor Protein (APP) affect å-endoproteolysis is only poorly understood. Thereby, mutations in the cleavage domain reduce å-efficiency of ã-secretase cleavage and some even shift entry into production lines. Since cleavage occurs within the TMD, a relationship between processing and TMD structure and dynamics seems obvious. Using molecular dynamic simulations, we dissect the dynamic features of wild type and seven FAD-mutants into local and global components. Mutations consistently enhance hydrogen8 bond fluctuations upstream of the ε-cleavage sites but maintain strong helicity there. Dynamic perturbation response scanning reveals that FAD-mutants target backbone motions utilized in the bound state. Those motions, obscured by large-scale motions in the pre-bound state, provide (i) a dynamic mechanism underlying the proposedcoupling between binding and å-cleavage, (ii) key sites consistent with experimentally determined docking sites, and (iii) the distinction between mutants and wild-type.

## Introduction

The onset and progression of Alzheimer’s disease (AD) are assumed to be linked to the accumulation of cell cytotoxic assemblies of β-amyloid (Aβ) peptides in the brain (Selkoe & Hardy, 2016). These Aβ-peptides are products of sequential proteolytic processing of the amyloid precursor protein (APP), an integral type I membrane protein (Haass, Kaether, Thinakaran, & Sisodia, 2012; Kaether, Haass, & Steiner, 2006). In the first step, ectodomain shedding of APP by β-secretase generates a 99 amino-acid long membrane-bound C-terminal fragment (C99). In subsequent steps, proteolysis of the C99 transmembrane domain (TMD) by the intramembrane-cleaving, aspartyl protease γ-secretase (GSEC) results in release of the APP-intracellular domain (AICD) and Aβ production (Lichtenthaler, Haass, & Steiner, 2011). Amyloidogenic processing of C99 (outlined in Figure 1) starts at the cytoplasmic border of the APP TMD at residue 48 or 49 (ε-sites) and progresses stepwise by three to four amino acids towards the N-terminus (Fukumori, Fluhrer, Steiner, & Haass, 2010; Matsumura et al., 2014; Olsson et al., 2014; Qi-Takahara et al., 2005; Quintero-Monzon et al., 2011; Takami et al., 2009). Depending on the initial ε-site, AICD49 or AICD50 and Aβ peptides of different lengths are released. In the wild-type (WT) form, the most abundant fragment is Aβ40, while Aβ42 and Aβ38 show significantly lower levels. Being highly prone to aggregation and a major component in senile plaques (Saito et al., 2011; Sandebring, Welander, Winblad, Graff, & Tjernberg, 2013) Aβ42 is probably the most neurotoxic species.

**Figure 1.**
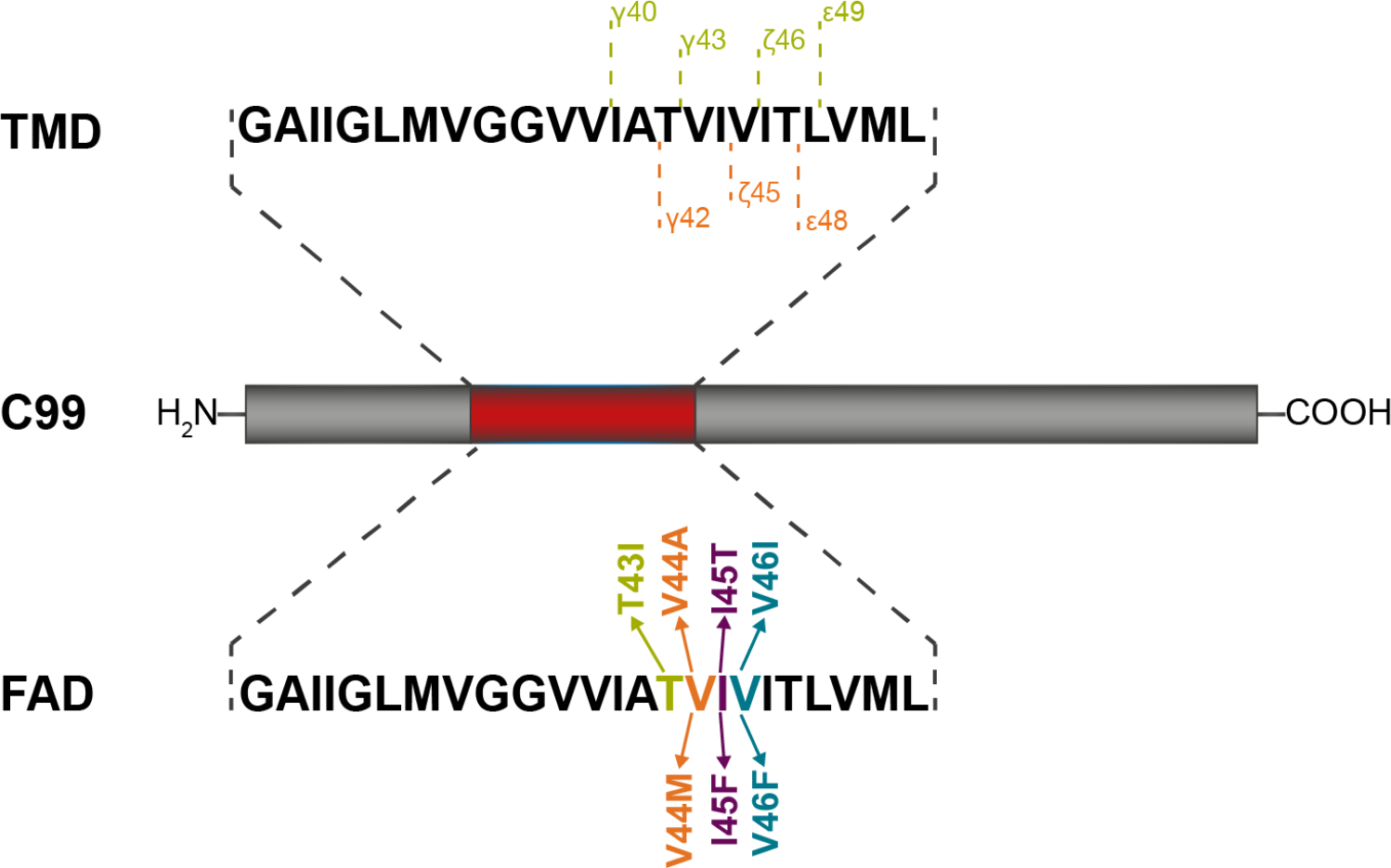
The C99 fragment of the amyloid precursor protein. The sequence of the transmembrane domain (residues G29 to L52) is shown in the upper panel together with the two main production lines which lead to formation of Aβ40 and Aβ42 peptides, respectively. The lower panel shows the location of FAD mutations investigated in this work.

Familial forms of AD (FAD), caused by mutations within the C99 TMD, commonly show major shifts in Aβ-ratios towards higher levels of Aβ42, where an increased Aβ42/Aβ40 ratio is correlated with the age of onset of AD (Alzforum, 2018; Chávez-Gutiérrez et al., 2012; Dimitrov et al., 2013; Haapasalo & Kovacs, 2011; Kakuda et al., 2006; Page et al., 2010; Richter et al., 2010; Weggen & Beher, 2012; Xu et al., 2016). Furthermore, most FAD-linked C99 mutant proteins are less efficiently cleaved by GSEC than WT as revealed by reduced generation of AICD and/or total Aβ levels (Bolduc, Montagna, Seghers, Wolfe, & Selkoe, 2016; Chávez-Gutiérrez et al., 2012; Xu et al., 2016). It should be noted that ε-efficiency is referred to as the proteolytic activity of GSEC (Quintero-Monzon et al., 2011). The changed Aβ42/Aβ40 ratio was related to a shift in preferential initial cleavage from the ε49-to the ε48-site (Qi-Takahara et al., 2005; Toru Sato et al., 2003; Takami et al., 2009). Recent findings, however, have demonstrated that final cleavage can be uncoupled from initial ε-cleavage and entry into a specific production line (Bolduc et al., 2016; Dimitrov et al., 2013; Matsumura et al., 2014; Olsson et al., 2014; Szaruga et al., 2017). Although the FAD mutants I45F and I45T enter the Aβ40 product line, both mutants increase the Aβ42/Aβ40 ratio. Thereby, the I45F-induced shift involves direct Aβ46-Aβ42 cleavage, skipping Aβ43 and Aβ40 production (Bolduc et al., 2016). V46F substitution drastically shifts ε-cleavage to ε49 (increased AICD50) and enters the Aβ42 product line. Nevertheless, it generates substantial amounts of Aβ40 (Dimitrov et al., 2013; Szaruga et al., 2017) indicating that it also used the alternative Aβ48-Aβ43-Aβ40 cleavage steps (Matsumura et al., 2014).

What determines initial ε-cleavage and ε-efficiency of C99? As cleavage occurs in the TMD of the substrate, the relevance of structural and dynamic features of the substrate’s TMD itself for processing seems obvious. An initial model assumed that the α-helical TMD must be locally destabilized at the cleavage site to be accessible for hydrolysis. However, biophysical analysis and *in-silico* modelling have revealed that the C-terminal cleavage region (TM-C) of C99 displays a strong α-helicity and is actually less flexible than the N-terminal region (TM-N) (Chen et al., 2014; Dominguez, Foster, Straub, & Thirumalai, 2016; Pester, Barrett, et al., 2013; Pester, Götz, Multhaup, Scharnagl, & Langosch, 2013; Takeshi Sato et al., 2009). Additionally, studies of threonine to valine mutations, located in the cleavage domain of the APP TMD did not show significant differences in ε-site flexibility and water accessibility (Scharnagl et al., 2014). Biochemical analyses have reported, however, that these mutations shift initial cleavage as well as cleavage efficiency and Aβ-ratios (Oestereich et al., 2015). Several NMR studies have reported differentially increased flexibility around the ε48-site in FAD mutants compared to WT (Chen et al., 2014; J.-X. Lu, Yau, & Tycko, 2011; Takeshi Sato et al., 2009). Some of these studies explicitly investigated the C99 TMD in the homo-dimeric state (Chen et al., 2014; Takeshi Sato et al., 2009), where contact interactions and increased C-terminal hydration might modify the properties of the C99 TMD (Pester, Barrett, et al., 2013). Recent studies, however, have revealed that APP in its dimeric state is not a substrate of GSEC (Fernandez et al., 2016; Winkler, Julius, Steiner, & Langosch, 2015).

Ultimately, substrate processing involves the subtle balance of interactions between substrate, membrane and enzyme in the crowded environment of the cell membrane. A certain degree of specificity may, therefore, be ascribed to the interactions of the substrate TMD with the enzyme. Hence, rather than switching ε-preference by locally destabilizing one ε-site over the other, FAD mutations might influence interactions and substrate processing prior to chemical cleavage. Models of enzymatic processing have provided evidence that the conformational plasticity of substrate and enzyme plays a key role for both recognition and subsequent relaxation steps (Haliloglu & Bahar, 2015; Henzler-Wildman et al., 2007; Ma & Nussinov, 2010; Wei, Xi, Nussinov, & Ma, 2016). Generally, functional dynamics results from the combination of intrinsic and induced effects: The intrinsic dynamics of the substrate prior to binding is essential to enabling large-scale cooperative changes in the structure, while induced motions, usually more localized, help to optimize and stabilize the bound conformations. At all steps, conformational flexibility of GSEC (Aguayo-Ortiz, Chávez-García, Straub, & Dominguez, 2017; Bai, Rajendra, Yang, Shi, & Scheres, 2015; Elad et al., 2015; Somavarapu & Kepp, 2016, 2017) might play a critical role. Furthermore, recognition, dynamics and conformation of the substrate TMD, as well as the enzyme, are influenced by the membrane environment (Barrett et al., 2012; Holmes, Paturi, Ye, Wolfe, & Selkoe, 2012; J.-X. Lu et al., 2011; Osenkowski, Ye, Wang, Wolfe, & Selkoe, 2008; Winkler et al., 2012).

Current models of substrate processing by GSEC (Figure 2) suggest initial binding of the substrate at an auxiliary interaction site (exosite) before it interacts with more distant recognition sites and finally gains access to the active site. The exosites are unknown, although Fukumori and Steiner showed that binding of C99 to GSEC is a two-stage process (Fukumori & Steiner, 2016). Their results further revealed that in the cleavage-competent complex, TM-N is in close contact with the N-terminal fragment (NTF) of presenilin-1 (PS1). In this complex, residues A42, V44, I45, L49, M51 and L52 in TM-C make contact with the docking site in the C-terminal fragment (CTF) of PS1. Recognition of TM-N has been discussed previously (Barrett et al., 2012; Fernandez et al., 2016; Kornilova, Bihel, Das, & Wolfe, 2005; Osenkowski et al., 2008; Yan, Xu, Melcher, & Xu, 2017) and is also supported by a recent structural model of GSEC complexed with a helical peptide (Bai et al., 2015). The site of initial ε-endoproteolysis was suggested to be dictated by residues upstream to the ε-site, i.e. I45 and V46 (Fernandez et al., 2016). Local unwinding of the scissile bond is coupled with entry of TM-C into the active site and binding the three residues downstream the cleavage site to the hydrophobic S1′-S2′-S3′ pocket within GSEC (Bolduc et al., 2016).

**Figure 2.**
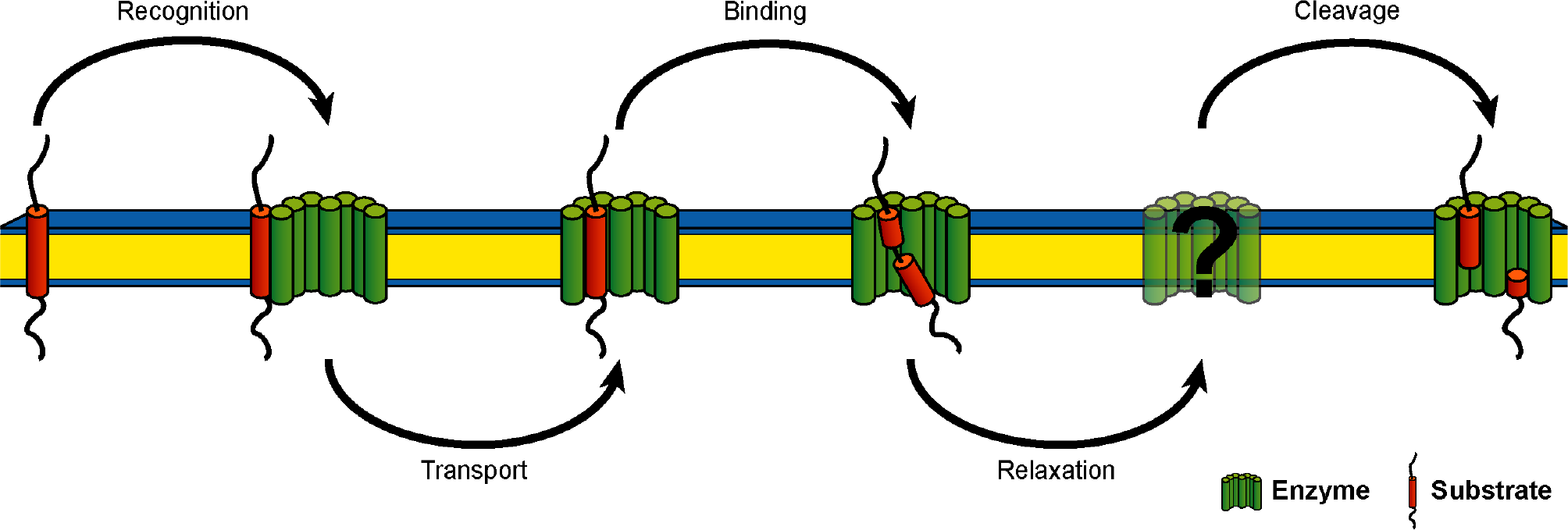
Schematic representation of catalytic processing of APP by γ-secretase. The five elementary steps represent 1) substrate recognition, 2) substrate transport to the docking site, 3) substrate binding and formation of the enzyme-substrate complex, 4) several steps of conformational reorganization of the enzyme-substrate complex, and 5) proteolysis and release of cleavage products. The proposed model is based on several mechanistic and structural studies of GSEC and APP as discussed by Langosch et al. (Langosch et al., 2015).

What characterizes the conformational plasticity of the C99 TMD and how can its dynamic properties contribute to binding and catalytic processing? NMR experiments and molecular dynamic (MD) simulations have suggested that the highly flexible G37G38 sites in the C99 TMD can act as dynamic hinges, where hinge-bending is triggered by the molecular environment (Barrett et al., 2012; Dominguez, Meredith, Straub, & Thirumalai, 2014; Lemmin, Dimitrov, Fraering, & Dal Peraro, 2014; Nadezhdin, Bocharova, Bocharov, & Arseniev, 2011). This di-glycine hinge has attracted a certain amount of interest because hinge-bending may assist a “swinging-in” of TM-C (Tian et al., 2002) to make contact with the GSEC’s active site. Changing interactions in TM-C by threonine to valine mutations were shown to modulate extent and direction of helix bending without altering local flexibility at the ε-sites (Scharnagl et al., 2014). The perturbed orientation of TM-C relative to the putative binding domain of TM-N provides an explanation for the reported shift in initial cleavage and cleavage efficiency (Oestereich et al., 2015). From this finding, an alternative view of ε-site selection was derived, relating altered cleavage site presentation to altered large-scale bending flexibility of the substrate TMD (Langosch, Scharnagl, Steiner, & Lemberg, 2015; Langosch & Steiner, 2016; Scharnagl et al., 2014; Stelzer, Scharnagl, Leurs, Rand, & Langosch, 2016). Notably, it was recently shown that cleavage of the α-helical SREBP-1 substrate by the intramembrane site-2 protease may also be associated with a hinge-like motion (Linser et al., 2015).

Based on our previous results (Pester, Barrett, et al., 2013; Pester, Götz, et al., 2013; Scharnagl et al., 2014; Stelzer et al., 2016), we suspected that FAD mutants may perturb the large-scale bending motion, which may in turn assist TM-C positioning in the enzyme’s active site (Langosch et al., 2015). Since global motions are enabled by local motions, FAD mutants are expected to induce changes in the helix-stabilizing network of backbone hydrogen-bonds (H-bonds). To probe our predictions, we studied the dynamics of model peptides of the C99-TMD for WT and seven FAD mutants (Figure 1) by molecular dynamics (MD) simulations. Due to the lack of a structural model of the substrate-enzyme complex, atomistic details concerning the nature of interactions are unknown. Hence, we used a matrix of low-dielectric, helix-stabilizing 2,2,2-trifluoroethanol (TFE) with low water content (80%/20% v/v) to mimic the interior of the enzyme (Buck, 1998). We showed that the TFE layer wrapping the peptide contains sufficient water molecules to satisfy hydration of hydrophilic sites, e.g. glycine and threonine. The TFE/H_2_O mixture thus provides a rational approach to the interior of GSEC, which captures significant amounts of water (C. Sato, Morohashi, Tomita, & Iwatsubo, 2006; Tolia, Chávez-Gutiérrez, & De Strooper, 2006). The simulations point out a substantial impact of the mutations on the stability of H-bonds in the cleavage region between γ-and ε-sites - however, neither H-bond flexibility nor water accessibility around the ε-sites showed significant differences. Rather, the alteration of the H-bond network fine-tunes the orientation of the scissile bonds relative to the putative TM-N binding region. However, this effect was unable to unambiguously distinguish between WT and mutants. Decomposition of the backbone dynamics revealed an extended motional repertoire complementing large-amplitude helix-bending. In order to gain insight into the functional role of this conformational diversity, we implemented a novel perturbation-response method. Here, the TMD in the TFE/H_2_O matrix defines a pre-bound state without specific interactions with the enzyme. Using a spring model, accounting for non-covalent substrate-enzyme interactions, we were able to scan the response of the TMD backbone dynamics to binding-induced perturbations. We found that tight packing of TM-N - as suggested by a recent hypothetic binding model (Bai et al., 2015) - obstructs large-scale helix bending in favour of motions localized in TM-C, which harbours the cleavage sites. This clearly translates into an altered presentation of the ε-sites in the FAD mutants as compared to WT. In the conformational ensemble of the pre-bound state, these bound-like motions are hidden. Their selective utilization in the bound state indicates that functional motions of the substrate TMD might differ from those large-amplitude motions contributing most to backbone flexibility. Our investigations propose a dynamic mechanism underlying the coupling between binding and presentation of the initial cleavage sites.

## Results

Here we used all-atom MD simulations to study the backbone dynamics of the C99 WT TMD and seven of its FAD mutants on a local and global scale. Properties were studied in a TFE/H_2_O mixture (80%/20% v/v) which favours the α-helical fold and satisfies both polar and hydrophobic interactions with the substrate TMD - an environment intended to mimic the interior of the enzyme (Buck, 1998). None of the investigated parameters indicates persistent conversion to an alternative backbone fold. Rather, they suggest the existence of a continuum of helical conformations, where helices bend, twist and straighten, but remain α-helical on average. This is well-documented by low root mean-squared deviations (RMSD) of the average structures with respect to an ideal α-helix which vary between 1.04 Å and 1.27 Å. RMSD values with respect to the average WT structure are even lower (RMSD_min_ = 0.15 Å for V44M and RMSD_max_ = 0.33 Å for T43I).

### FAD Mutations Induce Small Perturbations in Local Topology and Dynamics of the APP TMD

We systematically calculated and characterized local structural and dynamic differences of C99-TMD model peptides for WT and FAD mutants (for sequences see Methods and Figure 1). The analysed metrics covered structure (backbone dihedrals, rise-per residue), dynamics (Cα-fluctuations, occupancies of α- and 3_10_-H-bonds), and interactions (side-chain to main-chain packing, hydration). Deviations from the helical regime inform about backbone distortions as well as locally changed accessibility of the scissile bonds to hydrolysis. H-bond occupancies (Figure 3A) and root-mean squared fluctuations (RMSF) of Cα atoms (Figure 3B) revealed a heterogeneous distribution of backbone flexibility: A highly flexible centre is flanked by more rigid domains around G33 in the N-terminus (TM-N) and V46 in the cleavage domain (TM-C). In the flexible domain, transient breaking of H-bonds between the carbonyl oxygens from residues G33, L34, M35, V36 to the amide hydrogens of residues G37, V38, V39 and V40 could be observed. This is consistent with helix bending over the di-glycine hinge (Barrett et al., 2012; Pester, Barrett, et al., 2013; Scharnagl et al., 2014) where H-bonds on the convex side have to stretch in order to allow for bending. The flexibility imparted onto the helix is related to extensive shifting between α-helical and 3_10_-helical H-bonds (~25% non-canonical i, i+3 H-bonds) allowing the accommodation of TMD structural deformations (Cao & Bowie, 2012; Högel et al., 2018). Surprisingly, the FAD mutations investigated mainly targeted H-bonds located upstream of the mutation sites, i.e. in the region that harbours the γ and ζ cleavage sites. In contrast, FAD mutations had only a negligible impact at the ε-cleavage sites which showed preserved, stable H-bonding with >90% α-helicity. Remarkably, the H-bond pattern around the central di-glycine hinge was not or only slightly affected by the FAD mutations.

**Figure 3.**
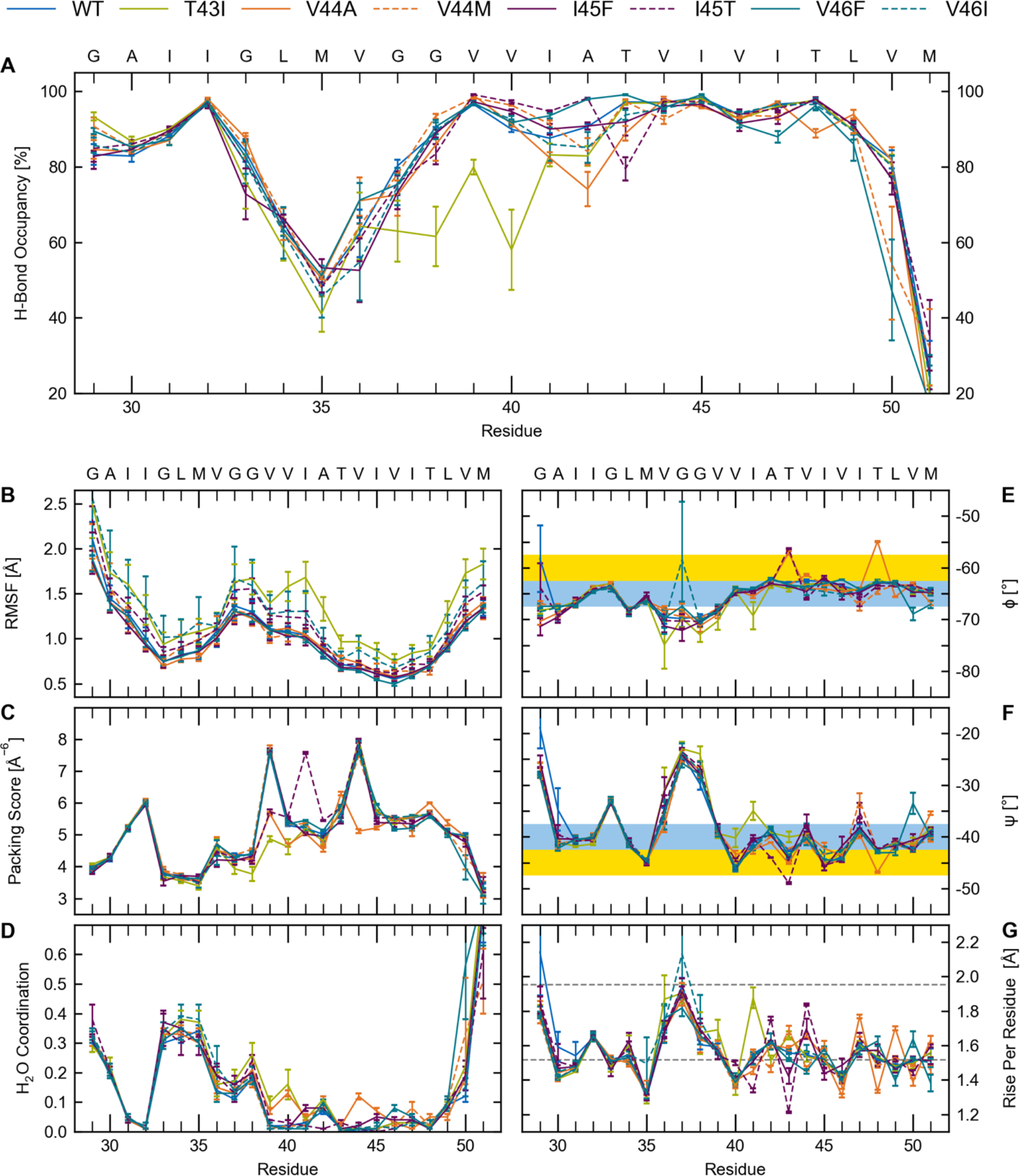
Structural and dynamic characterization of WT and FAD C99-TMDs. (**A**) Occupancies of intra-helical H-bonds between the carbonyl oxygen at position (i) and amide hydrogens at position i+4 or i+3. (**B**) Root mean-squared fluctuations (RMSF) of C_α_ atoms. (**C**) Packing of the carbonyl oxygen by side-chain atoms (values are multiplied by 100). (**D**) Number of water molecules within 2.6 Å around the backbone carbonyl oxygen. (**E, F**) Backbone dihedral angles. Blue areas indicate the torsion angle range of an ideal water-soluble helix, yellow areas indicate torsion angle range of ideal TM helices (**G**) Rise per residue between Cα atoms. Grey lines indicate values for ideal α-helices (1.52 Å) and 3_10_ helices (1.96 Å).

Packing scores (Figure 3C) are in close agreement with the H-bond occupancies. Side chains pack tightly around backbone carbonyl-oxygen atoms in TM-C, whereas packing deficiencies are present in the glycine-rich TM-N. The packing scores are mirrored by water coordination, which shows high values around glycine backbones and very low amounts of water molecules in the cleavage domain (Figure 3D). In the region between residues V39 and V44, the packing scores differ among WT, T43I, V44A and I45T. This can be related to perturbed back-bonding of the threonine side-chain. Depending on sequence context, back-bonding of a threonine side chain towards the protein’s main chain can induce both helix stabilizing and destabilizing effects (Deupi et al., 2010). Previous studies have revealed a rigidifying effect on the WT TMD due to side-chain to main-chain back-bonding of T43 and T48 (Pester, Barrett, et al., 2013; Scharnagl et al., 2014; Stelzer et al., 2016). For the FAD mutants studied here, analysis of the packing score (Figure 3C) indicates, that these helix-stabilizing interactions are perturbed in multiple ways. Besides the obvious lack of ability to bind back (T43I), both, I45T and V44A, drastically altered the native H-bond network. In contrast to our expectations, back-bonding of the threonine side chain of I45T to the I41 main chain does not provide additional stabilization when compared to WT. Instead, it forces the side chain of T43 to form a H-bond with its own backbone (80%). In the case of V44A, A44 hinders back-bonding of the T48 side chain and breaks back-bonding between T43 and V39. This forces the side chains of T43 and T48 to form alternative H-bonds either to their own backbone (~85%) or to the backbone carbonyl at the (i-1) site (~5%). The induced helix distortions are related to reduced H-bond occupancy (Figure 3A) at the T43 position in the I45T mutant, and at the A42 position (with a minor change at T48) in the V44A FAD mutant, respectively. Parallel to a reduced packing score, hydration increases at positions 40 and 44 (Figure 3D).

Finally, we discuss helix conformational distortions in terms of backbone dihedrals and rise per residue (Figure 3E-G). Generally, the backbone torsion angles of ideal transmembrane helices (Φ/Ψ ~ −60°/-45°) differ from their values for water-soluble helices (Φ/Ψ ~ −65°/-40°) (Kim & Cross, 2002). This 5° counter-directional change has a significant effect on the shielding of polar carbonyl groups from the surrounding environment (Page, Kim, & Cross, 2008). Accordingly, the Ψ values recorded for WT and FAD mutant TMDs indicate water-exposure of carbonyls G33, A42, V44 and I47, while the carbonyls at M35, V40 and I45 are more shielded. However, with one exception (M50 in the V45F mutant) torsions at the ε-sites do not point to increased water exposure. For G37 and G38 the strong negative Φ and the less negative Ψ translate to larger angles between adjacent peptide planes (Page et al., 2008). Thus, the carbonyl-oxygens are poorly H-bonded to their intra-helical partners but become more exposed to solvent. This is in accordance with H-bond occupancies (Figure 3A), water coordination (Figure 3D) and rise-per-residue (Figure 3G). Alternating backbone Ψ-torsions along the helices also translate into rise-per-residue values alternating between elongated and shortened Cα-Cα distances (Figure 3G) which should be 1.52 Å for ideal α-helices and 1.96 Å for 3_10_ helices (Guo, Kraka, & Cremer, 2013). Thereby, differences in the rise between residues flanking the peptide bonds of the two ε-cleavage sites are of particular interest. A significantly larger T48-L49 distance in combination with a shorter L49-M50 distance would indicate preference towards ε48 cleavage and formation of AICD49. The distributions of these distances for WT and FAD mutants (Figure 4) indicate a minor shift (~0.3 Å) towards increased T48-L49 rise-per-residue only for the V46F mutant. The shortened T48-L49 distance noted for the V44A mutant should even protect from ε48-cleavage.

**Figure 4.**
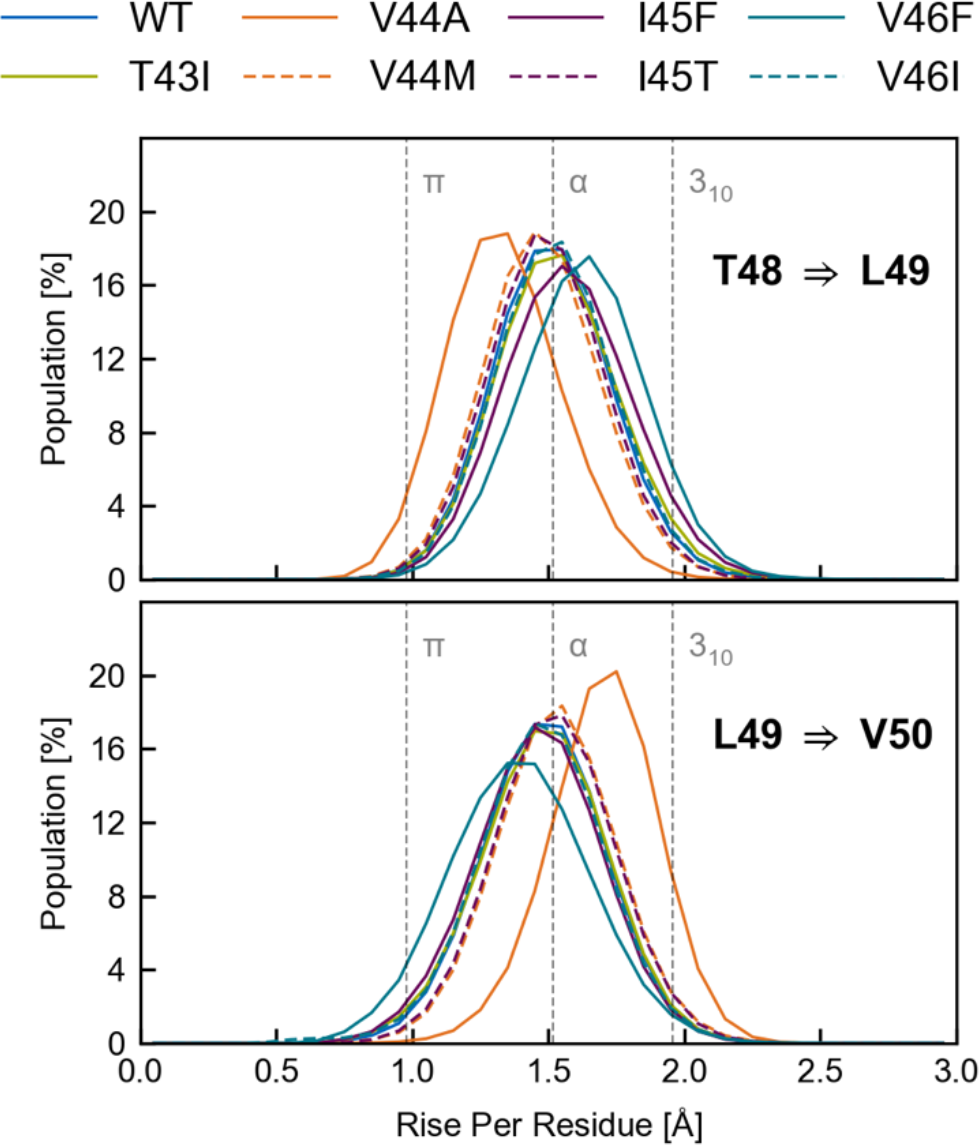
Rise between the residues flanking the scissile peptide bonds at the ε-sites. A shift in ε-preference towards ε48 cleavage would be indicated by a significant shift of the distribution of T48-L49 distances (upper panel) towards higher values as compared to the L49-M50 distances (lower panel). The bin width of the histogram is 0.1 Å.

In summary, the investigated parameters do not support a model where FAD mutants affect ε-cleavage due to local destabilization of an ε-site. Rather, they revealed an impact on H-bond flexibility in the region upstream to the ε-sites harbouring the γ and ζ cleavage sites.

### Local Fluctuations Cooperate to Generate Global Backbone Dynamics

Differences in the local structural dynamics alone cannot explain the impact of FAD mutations on ε-cleavage. Although localized at a single Cα-atom or H-bond, these fluctuations may cooperate to produce functional backbone flexibility enabling (i) large-scale conformational changes conducive to recognition and binding, (ii) induced conformational relaxations, and (iii) communication between functional sites distal in the sequence (Clarkson, Gilmore, Edgell, & Lee, 2006; Csermely, Palotai, & Nussinov, 2010; Ettayapuram Ramaprasad, Uddin, Casas-Finet, & Jacobs, 2017; Guarnera & Berezovsky, 2016; Haliloglu & Bahar, 2015; Marcos, Crehuet, & Bahar, 2011; Nashine, Hammes-Schiffer, & Benkovic, 2010). The dynamic cross-correlation mainly depends on the fold and is fine-tuned by sequence. Hence, the observed FAD mutation-induced variations within the H-bond network (see Figure 3A) may result in altered mechanical linkage properties directly affecting functionally important, correlated backbone motions.

The global backbone motions of a helical peptide can be intuitively related to the harmonic modes of a helical spring, namely elastic bending and twisting (Emberly, Mukhopadhyay, Wingreen, & Tang, 2003; Hub & de Groot, 2009; Pleiss & Jähnig, 1991) and are illustrated in **Supplementary Figure 5**. Previously, much attention has been paid to conformational changes of the APP-TMD related to hinge functionality of its central G37G38 motif (Barrett et al., 2012; Dominguez et al., 2014; Itkin et al., 2017; Langosch et al., 2015; Langosch & Steiner, 2016; Lemmin et al., 2014; Pester, Barrett, et al., 2013; Scharnagl et al., 2014). Hinges are flexible regions that permit the rotation of quasi-rigid segments around a screw axis passing through the flexible region. Accordingly, a helix that preserves its H-bonding structure when bent or twisted, provides mechanical hinges at the flexible sites (Hayward, 1999). The deformations associated with these changes are confined to a small number of hinge residues without altering the internal dynamics of the neighbouring quasi-rigid segments (Hayward, 1999). Here we aimed to identify regions with correlated low intra-segmental flexibility and to find the key structural motifs that control the inter-segmental motions of WT and FAD mutant TMDs. We compared structures sampled in the simulations with the mean structure using the Dyndom program (Hayward & Lee, 2002; Taylor, Cawley, & Hayward, 2014) (see Methods). Only a small percentage (< 5%) of the structures showed no deviations from a straight helix. As expected, motions of TM-N and TM-C helical segments coordinated by a single flexible region dominate (60-70% of sampled conformations, Figure 5A). The hinge is confined to 2-3 sites including the G37G38 motif (Figure 5B) and provides bending as well as twisting flexibility. In all peptides, bending significantly prevails over helix twisting (on average 45% bending, 20% twisting). Since the central G37G38 motif and its flanking valine residues in the APP-TMD do not feature packing constraints (see Figure 3C), they easily allow changes in the main-chain torsional angles (see Figure 3E-F) utilized to displace TM-C with respect to the TM-N region, as noted previously (Barrett et al., 2012; Dominguez et al., 2014; Lemmin et al., 2014; Scharnagl et al., 2014). For the APP-TMD, the sites with minimal RMSF (nodes of fundamental helix bending) are located at residues G33 and V46 (see also Figure 3B). According to a resulting distance of 13 residues, maximal bending of an ideal helix (**Supplementary Figure 5**) is expected to occur at sites V39/V40. The observed shift to G37/G38 (Figure 5C) in all C99 TMD peptides indicates the sequence-specific absence of packing constraints there. Unexpectedly, FAD mutations either maintain the hinge region or show a slightly higher preference for G38 (I45F, V46I, V44M) (Figure 5C). In ~20-30% of structures, shorter segments bend and twist around a pair of hinges localized in TM-N and TM-C - here referred to as “doublehinge” motion (Figure 5D). In the WT TMD, the two hinges are located at V36G37 and T43V44, respectively, and coordinate bending/twisting of the flanking segments relative to the middle domain. As a direct consequence of this coordinated motion, fluctuation changes in TM-N are coupled to fluctuation changes in TM-C. This results in a noticeable synchronous softening and shifting of the pair of hinges as result of FAD-induced H-bond loosening in TM-C.

**Figure 5.**
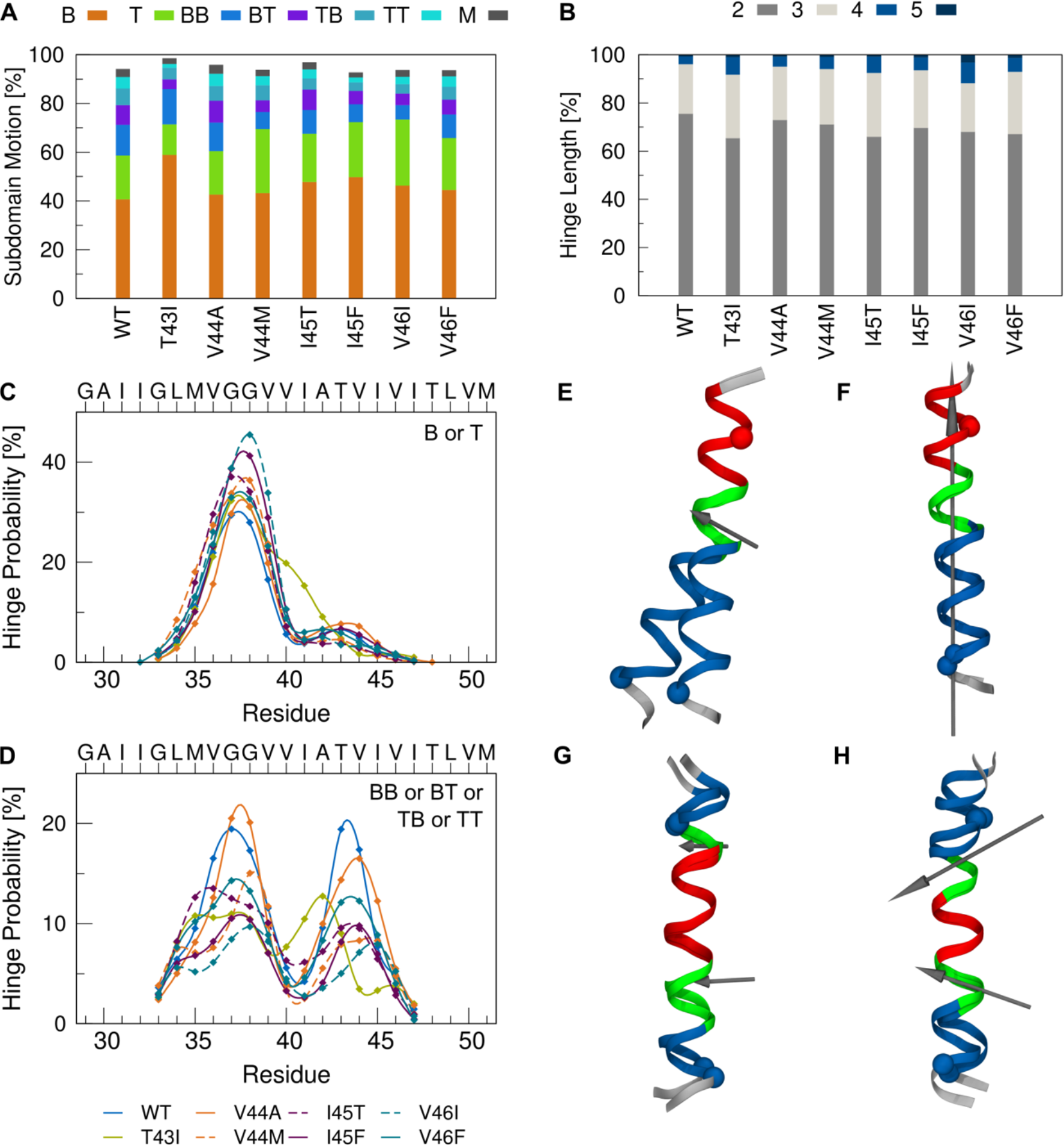
Decomposition of backbone dynamics. **(A)** Probability of hinge bending and twisting motions. Motions coordinated by a single flexible hinge are referred to as type Bend 1 (B) and Twist 1 (T), respectively. Bending and/or twisting around a pair of hinges is characterized by four combinations of bending (B) and twisting (T) (motion types BB, BT, TB, TT). Motions coordinated by more than 2 hinges are grouped in category M. (**B**) Number of hinge residues utilized for single-hinge motions. (**C, D**) Probability by which a residue is identified as a hinge site in the single-hinge (**C**) and double-hinge (**D**) bending and twisting motions. (**E-F**) Examples for bending (**E**) and twisting (**F**) around a single hinge. (**G-H**) Examples of correlated bending (**G**) and mixed bending/twisting (**H**) around a pair of hinges. Helical segments moving as rigid domains are coloured in blue and red, the hinge residues are highlighted in green. Screw axes are shown in grey. A larger projection of the screw axis with respect to the helix axis indicates a higher percentage of twisting. Spheres label the Cα atoms of G33 in TM-N and L49 in TM-C.

In summary, changes in global backbone motions can be substantiated by the mutation-induced alteration of the intra-helical H-bond network. The large-amplitude bending motion around the central G37G38 hinge contributes most to overall backbone flexibility and is largely conserved in FAD mutants. Perturbations of local interactions upstream to the ε-site provoke imprecise correlated helix bending around a pair of hinges localized in TM-C and TM-N. Due to its lower amplitude, this motion is largely hidden in the backbone RMSF profile of the C99 TMDs.

### The ε-cleavage Sites: Rigid, but Highly Mobile

How does the C99 TMD utilize its inherent flexibility for orientation of its ε-cleavage sites? How does the orientation profile differ between WT and FAD mutants? To answer these questions, we analysed the orientation variability of the helical segment harbouring the ε-sites (residues I47-M51) with respect to a helical segment in TM-N (residues A30-L34). We thus quantified the relative orientation is in terms of the bending (θ) and the swivel (Φ) angles between these two helical turns (for definitions see **Supplementary Figure 1**). It must be emphasised that these angles do not quantify the local bending at the di-glycine hinge. Rather, they are global descriptors for accumulated helix distortions between L34 and I47 (Figure 5).

**Figure 6.**
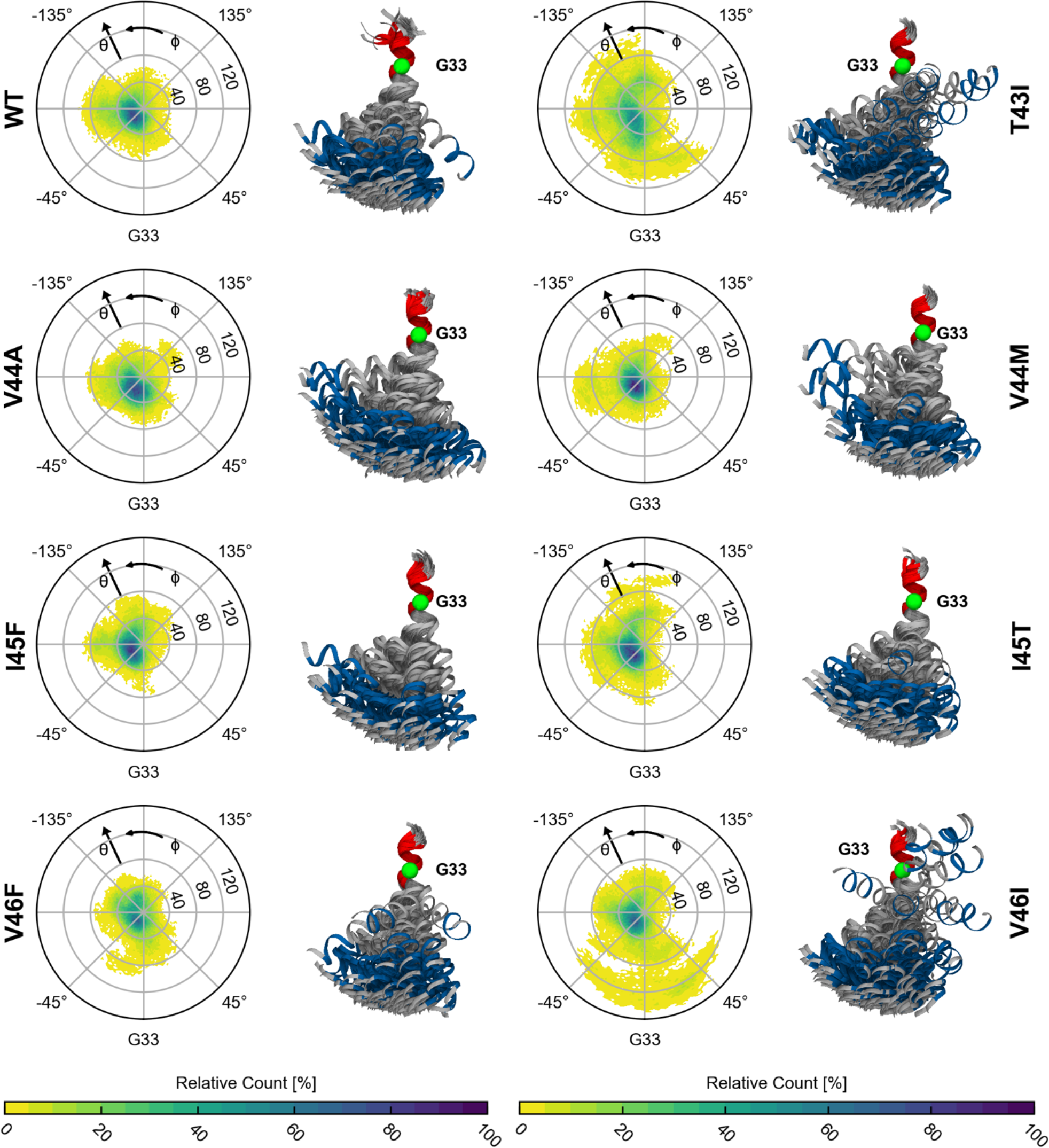
Cleavage domain orientation. Polar plots show 2D histograms of bending (θ) and swivel (Φ) angles characterizing the orientation of the helix segment I47-M51 carrying the ε-sites with respect to segment A30-L34. All values binned with a width of 2° were normalized to the bin with the overall highest count for all TMDs. Structures show an overlay of 150 frames with 1 ns spacing, visualizing the extent of shape fluctuations. Domains coloured in blue and red represent the selected reference segments for computation of angles. Green spheres represent Cα atoms of G33. The axis of the helical segment drawn in red has been aligned to the z-axis of the coordinate system and rotated around this axis to align the Cα atom of G33 with the x-axis.

As outlined above, the large-scale bending motion around the central di-glycine hinge is largely conserved in the FAD mutants (Figure 5C). Nevertheless, altered interactions and enhanced flexibility (see **Figure 3**) at residues C-terminal to the central G37G38 motif tune extent and direction of bending in the FAD mutants, mainly by loosening double-hinge bending (see Figure 5D). All TMDs mainly bend in direction of the di-glycine interface as depicted in Figure 6 and quantified by binning the conformations according to the swivel angle (Figure 7A). Note that for the coordinate system used in Figure 6 and **7**, G29, G33, G37 and G38 in an ideal α-helix are located at swivel angles Φ=40°, Φ=0°, Φ=-40°, and Φ=-140°, respectively. To further quantify differences between WT and mutants, conformations were classified in populations with small (0° ≤ *θ* < 20°), intermediate (20° ≤ *θ* < 40°) and large (*θ* ≥ 40°) bending angles (Figure 7B). In most cases, intermediate bending prevails. Conformations with large bending account for merely 10 to 20% of the total population, where T43I, I45T and V46I show a higher tendency towards larger bending angles combined with a larger variability of directions.

Like bending, stretching of the TMD might induce incorrect positioning of the ε-cleavage sites. To probe helix stretching, we compared the length of the helix axis connecting the ε-sites to G33 and G38, respectively. Here, the arc length of the helix between two sites is determined by the sum of the local rise parameters (see Figure 3G and Methods). For an ideally straight helix (rise per residue of 1.5 Å), the length G33→V50 (G33→L49) is ~25.5 Å (~24.0 Å), while the lengths between G38 and the ε-cut sites are expected to be ~7.5 Å shorter. With the exception of T43I, the peak values of the length distributions (Figure 7C) are only ~0.5 Å longer as predicted for a straight ideal α-helix. Variations from the ideal values mainly indicate helix stretching due to bending (Guo et al., 2013). While the extension of the C-terminal helix length reflects FAD-induced perturbations upstream of the ε-sites, the helix length measured up to G33 also includes helix distortions around the di-glycine hinge. The small populations with a ~4.5 Å longer helix axis observed for T43I and V46I, (Figure 7C, left panel) can be attributed to conformations where bending angles > 80° (see also Figure 6) drastically elongate the region around the di-glycine hinge.

To summarize, the orientation of ε-sites with respect to the N-terminal part of the C99 TMD is dominated by the signature of anisotropic bending over the G37G38 hinge and remains largely conserved in FAD mutants. Variability of the ε-presentation emerges from FAD-induced loosening of the double-hinge, contributing ~20-30% to the cumulative effect of all bending and twisting motions. Moreover, the analysis reveals that the ε-sites, while locally rigid, are highly mobile.

**Figure 7.**
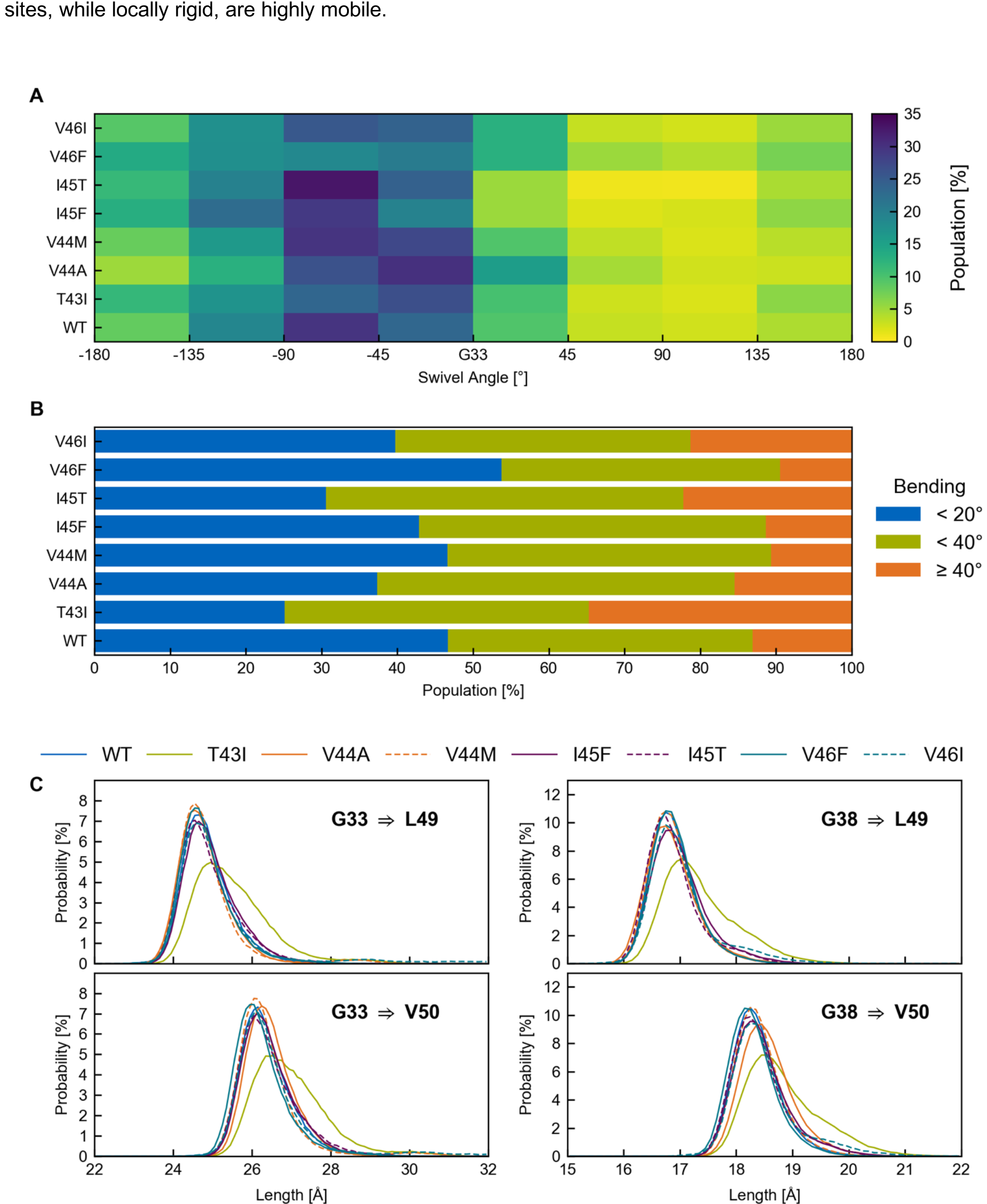
Statistics for swivel angles, bending angles and distributions of helix lengths. (**A**) Classification by swivel angle (Φ) in bins with bin size of 45°. The highest populations for all investigated peptides are found for orientations related to bending over the hinge (0° < Φ < −90° counted from G33). (**B**) Classification of conformations in populations with small (0° ≤ θ < 20°), intermediate (20° ≤ θ < 40°) and large (θ > 40°) bending (θ) angles. (**C**) Length distributions of accumulated rise per residues from the ε-sites to G33 and G38, respectively. The bin width of the histogram is 0.1 Å.

### Functional Backbone Motions Deduced from Impact of FAD Mutations on H-Bond Flexibility

Our previous results revealed that the ε-sites participate in a tightly coupled network of H-bonds. They remain locally rigid but gain mobility as a consequence of large-amplitude, collective backbone motions. How can we figure out the functional relevance of these motions for catalysis? Here we rationalize, that backbone motions which are targeted by FAD mutations in the C99 TMD, are of functional importance. The pronounced impact of FAD mutations on H-bonding makes the H-bond occupancies (Figure 3A) promising candidates for functional order parameters. We focused our analysis on the summary score for H-bonds emanating from carbonly oxygens in two regions, namely (i) from G33 to G38, and (ii) from V40 to T43. The backbone motions correlating maximally with these occupancy fluctuations (Hub & de Groot, 2009; Krivobokova, Briones, Hub, Munk, & De Groot, 2012) are termed type I (FM I) and type II (FM II) functional modes respectively. For model building and cross-validation see **Methods** and **Supplementary Figure 3**.

The observed backbone motions reveal large differences between mutants and WT as well as among the mutants. In general, the functional modes describe hinge bending and twisting motions of helical segments around various flexible joints, as identified with the Dyndom program (Hayward & Lee, 2002). The contributions of each residue to these motions vary largely (see Figure 8). For all peptides, motions correlating with H-bond fluctuations in the central region (type I) describe large-scale hinge bending localized in the residue segment from V36 to V39 (Figure 8A). However, even minor changes in H-bonding at this location can bias direction of bending or enhance twisting flexibility in type I motions. Accordingly, the bending-over-the-hinge pattern dominating the overall distribution of swivel angles (Figure 7A) is recovered only for a subset of peptides (WT, V44A, V44M and I45T), while the FAD mutants I45F, T43I, V46F and V46I bend in perpendicular directions and separate in different clusters (Figure 8B). The backbone motion enabled by H-bond fluctuations around the γ-site (type II) is a double-hinge motion in WT but shows a repertoire of bending and twisting around diverse hinge sites for the FAD mutants (Figure 8C). As a result, FM II of FAD mutants display the largest dissimilarity to WT as well as among the different mutants (Figure 8D and **Supplementary Figure 8-1**).

To summarize, backbone motions correlating with local H-bond fluctuations targeted by FAD mutations deviate from those large-scale motions contributing most to overall backbone flexibility. Strikingly, the variation between low and high H-bond occupancies in the two regions causes a substantial change in the directions of ε-site presentation.

**Figure 8.**
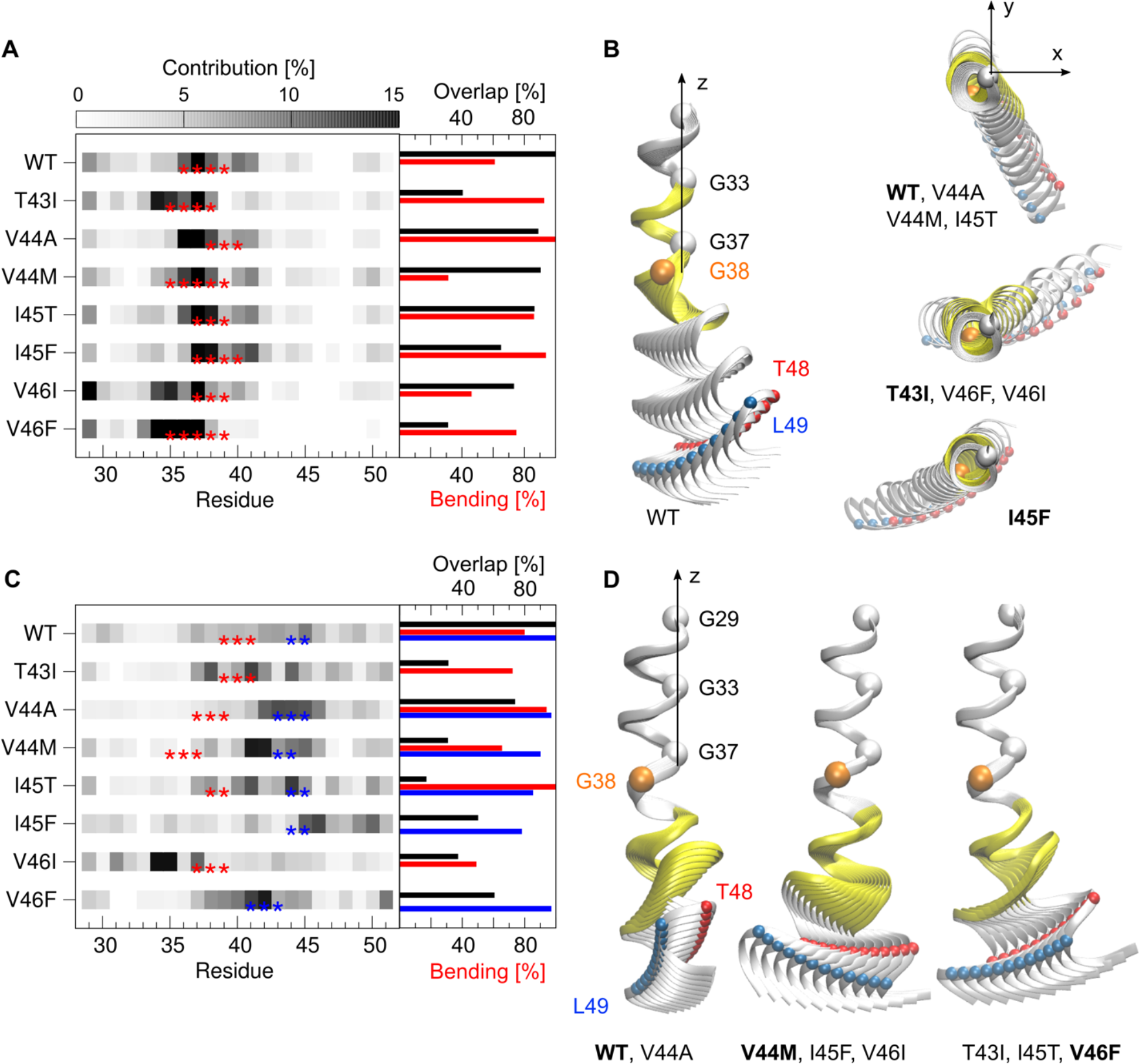
Functional mode analysis. (**A**) Amino acids contributing to the motions correlated with occupancy variations of H-bonds spanning residues G33-V42. Red stars indicate hinge residues. The percentage of helix bending is shown with red bars in the right part (%-twisting amounts to 100 - %-bending). Black bars quantify similarities between mutants and WT. (**B**) Backbone representation of structures interpolating from low to high occupancies along the motions from (**A**). (**C**) The same as for (**A**) for H-bonds spanning residues V40-I47. Hinge residues are highlighted by red and blue stars; percentages of helix bending are shown in accordingly coloured bars. (**D**) Backbone representation of structures interpolating from low to high occupancies along the motions from (**C**). For the graphical representations, structures have been overlaid onto the A30-L34 segment. The part of the backbone spanned by the corresponding H-bonds is coloured in yellow. Functional modes were clustered, the depicted cluster representative is indicated with bold letters. ε-cleavage sites (T48, L49) are shown as red and blue spheres, respectively.

### Functional Backbone Motions Deduced from Response to Binding-Induced Interactions

Surprisingly, backbone motions correlating with H-bond fluctuations targeted by FAD mutations appear to be mainly hidden in the overall backbone dynamics, which is dominated by large-amplitude helix bending. They might, however, be selectively utilized for optimization and relaxation steps after substrate-enzyme binding, where large-scale bending motions are obstructed by a tightly packed environment (Lezon & Bahar, 2010; Ma & Nussinov, 2010; Marcos et al., 2011). No structural model for the substrate-enzyme complex is currently available, though the experimentally determined structure of GSEC complexed with a co-purified α-helical peptide (Bai et al., 2015) provides an initial hypothetical model. In this case, PS1-NTF tightly embraces the N-terminal segment of the helix and provides various hydrophobic and polar contacts (see Figure 9A). The functional relevance of this distinct environment is supported by the high amount of PS1 disease mutations located there (Alzforum, 2018). Although lacking further atomistic details of the nature of interactions between substrate TMD and GSEC, our MD simulations allow the response of the TMD backbone dynamics to binding-induced perturbations to be studied. Here, we treat non-covalent interactions between substrate TMD and enzyme implicitly i.e. as a source of elastic energy. These interactions influence the motions of the explicitly treated mechanical degrees of freedom, which are represented by the Cα atoms of the helical core. Given the fluctuations in the pre-bound state, the reorganization of conformational dynamics can be probed for a variety of model interactions. Similar perturbation-based analyses have been employed in various previous studies (Ettayapuram Ramaprasad et al., 2017; Guarnera & Berezovsky, 2016; Ming & Wall, 2006; Zheng & Brooks, 2005; Zheng, Liao, Brooks, & Doniach, 2007). For further details and comparison with related approaches see Methods.

In the first step, we scanned fluctuation changes of each residue when binding occurs at each other residue of the WT-TMD. Figure 9B reveals that binding in one part of the TMD induces strong changes in the other part far from the binding site. This communication can be easily understood, since propagation of the binding perturbation depends on the collective correlation of residue fluctuations. Therefore, the resulting conformational changes are global, even in the event of local perturbations at one specific site. For example, binding at G33 significantly increases fluctuations at V44/I45. Thereby, the effect shows a pronounced nonreciprocity, i.e. binding at TM-C (e.g. V44 or T48) enhances flexibility at V36, while binding at V36 will not produce significant fluctuation changes in TM-C. This non-reciprocity is a direct consequence of a site’s acceptance for perturbation depending on its local flexibility. In the second step, we discussed simultaneous binding of multiple sites of the substrate WT-TMD to the enzyme, namely contacts with G37 and G38 (model 1), G33 and G37 (model 2), G38 and A42 (model 3), M35 and V39 (model 4), and residues from G29 to G38 (model 5). The G33xxxG37 and G38xxxA42 motifs were previously discussed as putative interfaces for C99 TMD homodimerization (Dominguez et al., 2014; Munter et al., 2007; Wang et al., 2011) and might also be involved in contacts with PS1 helices. M35 and V39 in model (4) are located on the opposite side of these motifs. Model 5 is based on the hypothetical binding model (Bai et al., 2015), where the helical segment incorporated within GSEC is 9 residues long, corresponding to residues G29 to G38 in our model. Depending on the interaction sites, binding-induced stiffening propagates along different pathways as reflected in a diverse pattern of fluctuation differences (Figure 9C). They range from minor communication (model 1) to increased fluctuations in TM-N, accompanied by decreased fluctuations in TM-C (models 2 and 3) up to above-average fluctuations in both, TM-N and TM-C (model 4).

**Figure 9.**
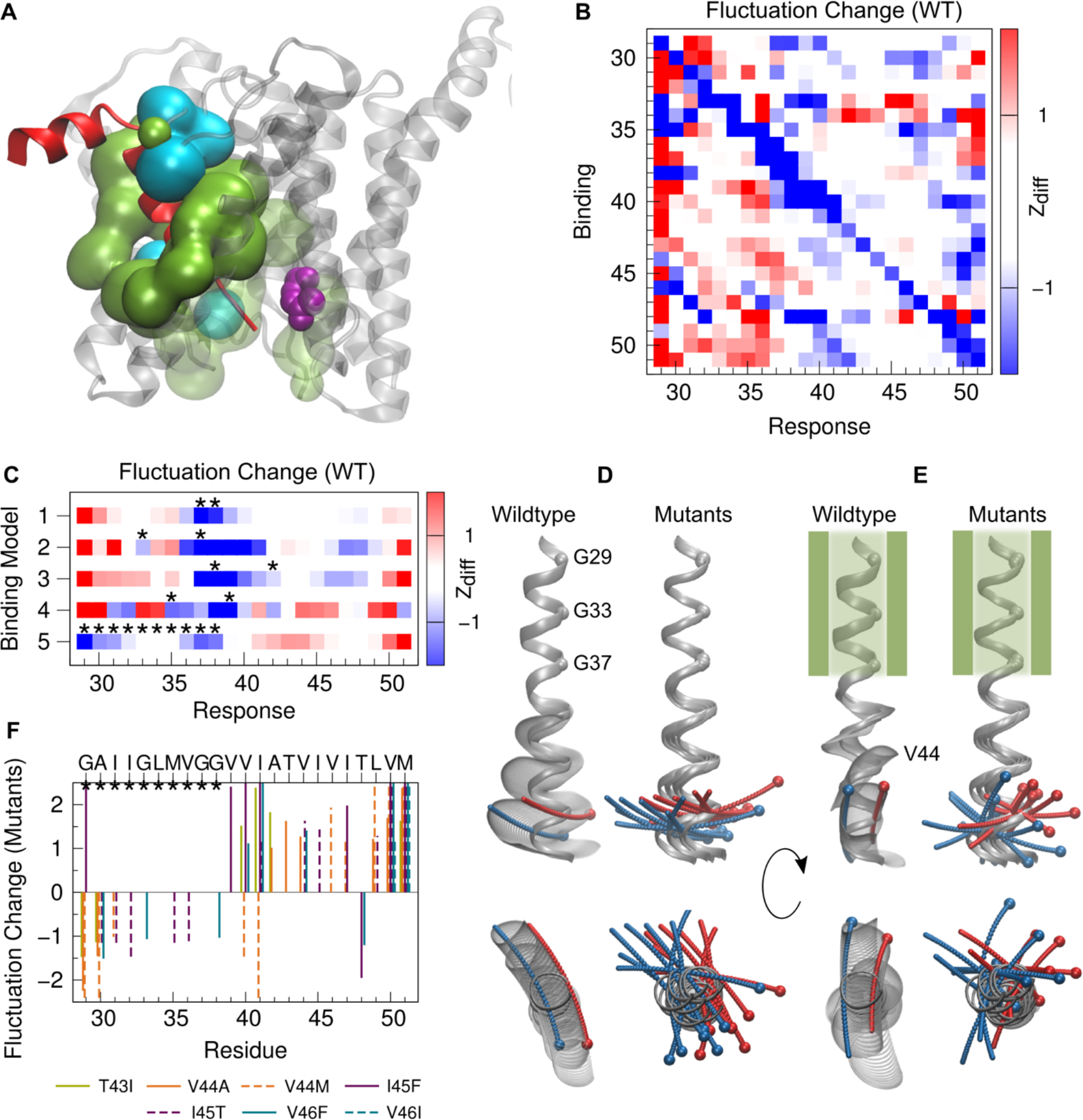
Response of TMD dynamics to binding interactions. (**A**) A hypothesis for substrate binding (Bai et al., 2015) (pdb 5FN3). PS1 (grey ribbons) residues within 7 Å distance to the bound helix (red) are shown in surface representation (green: hydrophobic residues, cyan: polar residues, residues contacting the unresolved part of the helix are drawn in transparent mode). The active site aspartates are shown in purple. (**B**) Perturbation response scanning for WT TMD. Z_diff_ quantifies the difference between normalized MSF profiles of bound and pre-bound TMD when binding is at a single residue. (**C**) Multiple site binding models for the WT TMD. Binding sites are indicated by black stars, Z_diff_ as in (**B)**. (**D, E**) Backbone motion of WT TMD (left) and backbone traces of the mean structures of FAD mutant TMDs (right) in the pre-bound (**D**) and bound state as defined by model 5 (**E**), respectively. For the graphical representations, structures have been overlaid to the segment A30-L34. ε-cleavage sites (T48, L49) are shown as red and blue spheres, respectively. (**F**) Significant Z_diff_ scores of normalized MSF differences between FAD mutants and WT for binding model 5.

In the following section, we compare the conformational ensembles in detail for binding model 5. As expected, large-scale bending, dominating the mode space in the pre-bound state (Figure 9D), is obstructed in the tight environment of the bound substrate TMD. By contrast, lower-amplitude motions - previously hidden in the global dynamics of the pre-bound state - are now selectively utilized (Figure 9E). In the case of the WT, reorganization of the TMD’s conformational dynamics in the bound state is exemplified in **Supplementary Figure 9-1**. The dominant modes after binding (Figure 9E) now favour local bending around residues A42-V44. Most importantly, binding sites in TM-N coupling to the same conformational degree of freedom (Figure 5D) can establish a communication pathway to TM-C, i.e. G33 and L34 can act as effectors, while A42-V46 act as sensors (see Figure 9B). Disease-associated mutations perturbing local flexibility and shifting hinge centres at the sensory sites (see Figure 5D) may worsen dynamic communication between TM-N and TM-C and impair precise positioning of ε-sites. The binding-induced shift of dominant backbone modes from central large-scale to more localized bending is reflected in the change of local mean-squared fluctuations (MSF). As shown in Figure 9C for the WT, fluctuations around G37G38 are reduced, but concomitantly increased around I41-I45. Significant fluctuation changes between FAD mutants and WT occur along the complete TM-C (Figure 9F). All mutants preserve low MSF at the ε48 site; I45F and V46F even reduce fluctuations there, while V44M and I45T respond with increased fluctuations at L49. Notably, all FAD mutations increase fluctuations at the carboxy-terminus (at residues V50 and M51).

The question that arises is how the previously defined functional modes compare to motions contributing most to overall backbone flexibility in the pre-bound and bound state? To clarify this question, we calculated the overlap of the functional modes (see Figure 8) with the essential dynamic subspaces describing 85% of the respective overall Cα MSF in both states. The functional modes overlap only poorly in the pre-bound state (Figure 10A). The overlap of type I motions is ~60% for the WT TMD and varies largely (σ=15%) around that value for the FAD mutants. Type II motions show consistently an even lower overlap (μ=~40%, σ=5%). In contrast to the poor overlap in the pre-bound states, the overlap of both types of functional modes with bound state essential motions is drastically enhanced (Figure 10B) and reaches >90% (FM I) and 74%-87% (FM II), respectively. This indicates that H-bonds which are affected by FADs and covered by FM II, are of significant importance to relaxations in the enzyme-substrate complex.

**Figure 10.**
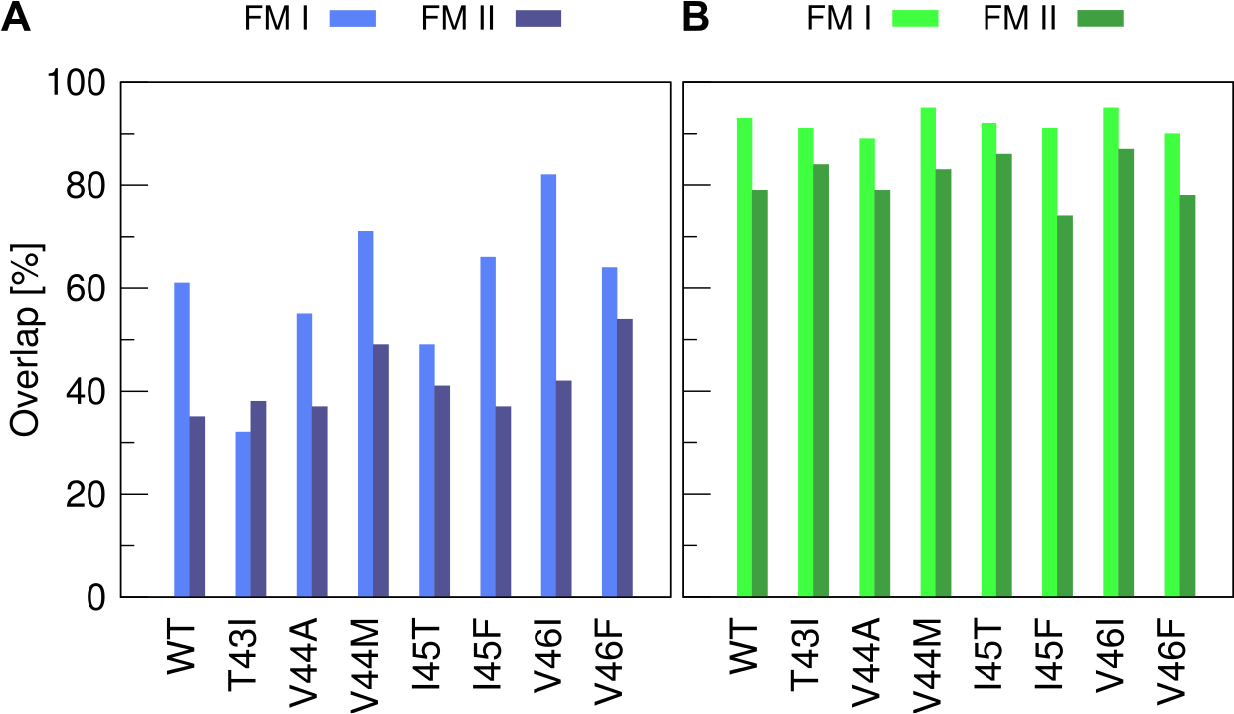
Similarity of functional modes and essential backbone motions. (**A**) Overlap with the essential subspace of the pre-bound TMDs. (**B**) Overlap with the essential subspace in the bound state where TMD dynamics is perturbed according to binding model 5. Functional modes of type I (FM I) correlate maximally with fluctuations of H-bonds spanning residues G33-V42. Functional modes of type II (FM II) correlate maximally with fluctuations of H-bonds spanning residues V40-I47.

To summarize, the response of conformational dynamics to binding interactions reveals remarkable C99-TMD plasticity. Instead of bending over the di-glycine hinge, a bound substrate TMD utilizes bending of terminal helix segments relative to the middle domain. In the pre-bound state, this motion is largely hidden and contributes only ~20-30% to overall backbone dynamics. Along this motion, binding-induced stiffening of the N-terminal part is communicated to the cleavage region supporting a model where initial interaction with the enzyme sets up the endoproteolysis site (Fernandez et al., 2016). The identified sensory sites in TM-C overlap with experimentally determined docking sites interacting with PS1-CTF (Fukumori & Steiner, 2016). Loosening or shifting the sensory sites in FAD mutants might compromise precise communication and presentation of the ε-cleavage sites.

### The TFE/Water Mixture: A Mimetic for the Interior of the Enzyme

2,2,2-trifluoroethanol (TFE) is perhaps the most commonly used agent for stabilization of α-helical conformations. In a mixture with water, this solvent can provide a variety of non-specific environments ranging from those mimicking hydrophobic interactions in the interior of proteins to hydrophilic ones at the surface (Buck, 1998; Roccatano, Colombo, Fioroni, & Mark, 2002). Here we mimic features of the substrate TMD in the interior of the GSEC enzyme using a TFE matrix containing a low amount of water (80%/20% v/v) to account for the presence of functionally important water molecules (X. Li et al., 2013; P. Lu et al., 2014; C. Sato et al., 2006; Tolia et al., 2006). What does this environment look like in our experiments and how might it be compared to the “real” enzyme? To answer those questions, we calculated the spatial distribution of TFE and water molecules around the WT TMD. While water enriches around the charged termini, the core (residues A30-V50) is preferentially solvated by TFE (Figure 11A). TFE clusters around hydrophobic residues in TM-C and TM-N with lower preference for the central G37G38V39 region (Figure 11B). The TFE layer still contains enough water molecules to satisfy hydration of hydrophilic residues (G29, G33, G38, T43). This observation is consistent with previously determined water coordination (Figure 3D). The increased local TFE concentration and shielding from water makes the environment of residues in TM-C (V40-L49) more apolar, thus stabilizing intra-helical polar interactions (backbone H-bonds and threonine back-bonding, compare also Figure 3A-C). The distributions of water and TFE molecules are visualized in Figure 11C by density surfaces. Residues in TM-C are in close contact with the hydrophobic part of TFE. The exposure of the TFE-hydroxyl group to the outer water phase indicates that TFE does not compete directly for H-bonding with the peptide backbone, i.e. the accumulation of TFE at the surface of the peptide is not caused by preferential H-bonding between the backbone and TFE. Around the mid-peptide region (G37G38), water penetrates closer. We conclude that the preferential association of the solvent with the peptide surface resembles a substrate TMD globally fitting into a compatible environment in the enzyme. The TFE layer provides a tightly packed apolar environment facing hydrophobic side chains, but also allowing water molecules to assemble around polar groups. In the real enzyme, polar contacts around hydrophilic side chains in the N-terminal region of the helix might be supplied by a set of polar residues (i.e. threonine and serine, see Figure 9A). Specific interactions, however, such as those provided by the catalytic aspartates, are missing from the solvent model.

**Figure 11.**
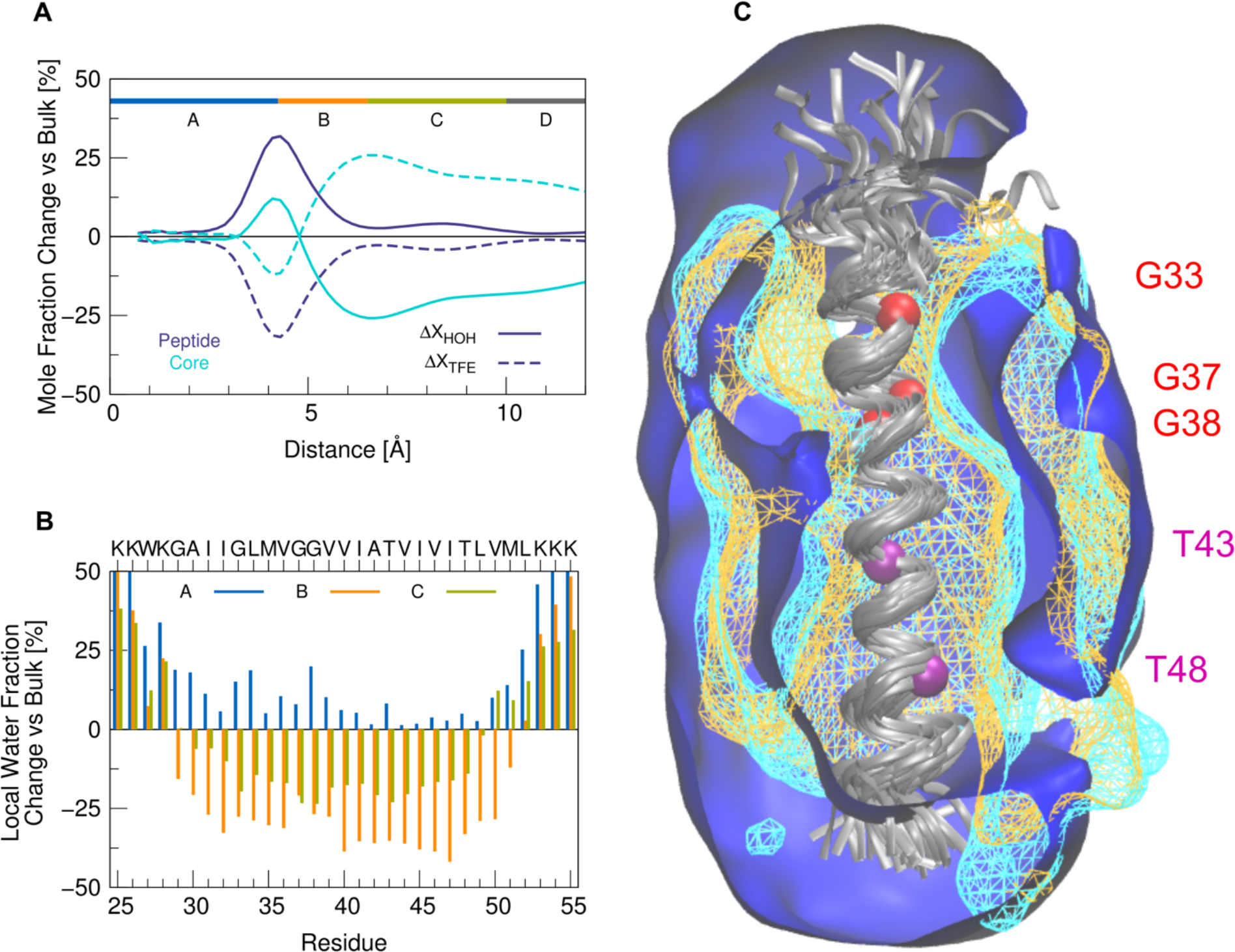
Distribution of solvent around the WT peptide in TFE/water. **(A)** Mole fraction of water (full lines) and TFE molecules (dashed lines) according to their distance from the peptide. The change is calculated with respect to the bulk values (X_HOH_ (bulk) = X_TFE_ (bulk) = 0.5). Compared is the distribution including all residues (dark blue) with the distribution around the hydrophobic core only (residues A30-V50, light blue). (**B)** Residue-specific change of the water fraction in three regions (A: r≤4.5 Å, blue; B: 4.5 <r≤ 6.5 Å, orange; C: 6.5 <r≤ 10 Å, green). **(C)** Contour plots of water (blue) and TFE (orange: hydrophobic part, cyan: hydroxyl group) drawn at a density of 2.3 10^−4^ molecules/Å^3^ corresponding to 35% bulk molarity. Data from the last 50 nanoseconds of the simulations were accumulated. The backbone of the peptide (grey) was superimposed onto residues G29-M51 of the average structure. Cα atoms of glycine and threonine residues are indicated by spheres in red and purple, respectively.

## Discussion

It is well understood that enzyme catalysis takes advantage of the repertoire of motions inherent to native proteins, although their role in the chemical step seems less obvious (Pisliakov, Cao, Kamerlin, & Warshel, 2009; Warshel & Bora, 2016). Transitions along large-scale motions might be selected to enable recognition, while lower-amplitude, localized motions help to optimize and stabilize enzyme-bound states (Boehr, Nussinov, & Wright, 2009; Haliloglu & Bahar, 2015; Henzler-Wildman et al., 2007; Ma & Nussinov, 2010; Nashine et al., 2010; Wei et al., 2016). Currently, no structural model for a GSEC-bound substrate is available and hence atomistic details concerning the nature of interactions and conformational relaxations involved in binding and presentation of the ε-site are unknown. Here we treated TMD-enzyme interactions in an implicit way and compared local and global dynamical properties of the TMDs of C99 WT and seven early-onset FAD mutations (T43I, V44(A,M), I45(T,F) and V46(I,F)). The chosen solvent (80%TFE/20% water v/v), intended to mimic for the interior of the enzyme, provides a non-specific environment that satisfies both, polar and hydrophobic interactions with the substrate TMD. Equilibrium dynamics of the substrate TMDs is calculated using all-atom MD simulations. The simulations have previously been validated by comparing MD-derived amide deuterium-hydrogen exchange rates with experiments (Pester, Barrett, et al., 2013; Pester, Götz, et al., 2013).

Mutations do not imply substantial changes in the mean structure, rather they affect the dynamic ensemble of conformations where the helices bend, twist and straighten, but remain α-helical on average. The FAD mutations consistently increase H-bond fluctuations in the region upstream to the ε-site while maintaining strong α-helicity near the cleavage sites. A stable helix near scissile bonds was found previously not only for the C99-TMD (Chen et al., 2014; Dominguez et al., 2016; Pester, Barrett, et al., 2013; Pester, Götz, et al., 2013; Takeshi Sato et al., 2009; Scharnagl et al., 2014), but also for Notch (Deatherage et al., 2017) and the SREBP-1 substrate of the intramembrane site-2 protease (Linser et al., 2015). The vicinity of the γ-site was shown, by solid-state NMR, to exist in a mixture of helical and non-helical conformations (Itkin et al., 2017; J.-X. Lu et al., 2011), consistent with reduced H-bond occupancies as revealed from our simulations. Significant differences within cooperative correlations emerge from these subtle differences within the underlying H-bond network. However, large-scale, global helix bending - primarily defined by the α-helical fold - is nearly unaffected by FAD mutations. Rather, the mutation-induced loosening of a pair of hinges in TM-N and TM-C was found to be a main determinant for orientation of the ε-sites. This double-hinge motion coordinates bending and twisting of the N-and C-terminal helical segments around the middle domain, with hinge-sites localized in TM-C around A42-I45 and in TM-N around V36-G38.

To gain insight into the functional importance of these backbone motions, we investigated how energetic perturbations, associated with enzyme interactions, cause and affect the reorganization of the TMD’s conformational dynamics. In our *in-silico* modelling study, we treated TMD-enzyme interactions implicitly as a source of elastic energy, weakly and harmonically restraining backbone atoms at binding sites. The propagation of the perturbation throughout the protein is mediated by the correlation of residue fluctuations in the pre-bound state. Introducing the quasi-harmonic covariance matrix determined from MD simulations as response function, we extended previous models based on harmonic formulation of protein dynamics (Erman, 2006; Guarnera & Berezovsky, 2016; Ming & Wall, 2005b; Zheng & Brooks, 2005; Zheng et al., 2007). In particular, the conformational reorganization depends on all protein backbone motions and is not restricted to a few so-called essential modes (Amadei, Linssen, & Berendsen, 1993). Thermodynamically, the redistribution of the TMD’s dynamic fluctuations entails changes in conformational entropy. The decrease in entropy due to binding-induced stiffening at contact sites might be compensated by an enthalpy gain due to new TMD-enzyme interactions and by the release of water from the binding interface. It can also be balanced by an increase in motion at other regions, thereby providing a means of propagating binding information to distal sites, even in the absence of significant structural changes. Highly flexible binding interfaces might, therefore, enhance functional important communication with sensory sites (Freire, 1999; Liu, Whitten, & Hilser, 2006; Pan, Lee, & Hilser, 2000), suggesting a special role for flexibility-enhancing glycine residues in TMDs (Högel et al., 2018).

Our perturbation-response approach allows us to scan a variety of binding models. One model was studied in greater detail. Based on recent structural data (Bai et al., 2015), we speculated that a hydrophobic interaction with PS1-NTF holds the TM-N of the C99 TMD and aligns it for catalysis. Our analysis reveals the importance of the region upstream to the ε-sites for conformational relaxation after binding. TMD dynamics in this bound state selectively utilizes the double-hinge motion, which contributes only ~20-30% to conformational variability in the pre-bound state. In exchange, the predominant bending motion in the pre-bound state is mostly obstructed in the bound state. Furthermore, the utilized conformational degree of freedom provides communication between binding to TM-N and presentation of the ε-sites, thus supporting a model, where initial interactions set up the site of endoproteolysis (Chávez-Gutiérrez et al., 2012; Fernandez et al., 2016) by GSEC. The identified sensory sites in TM-C overlap with experimentally determined docking sites interacting with PS1-CTF (Fukumori & Steiner, 2016). Loosening and/or shifting the sensory sites in disease-mutants might compromise precise presentation of the ε-sites to the active-site cleft in the enzyme-substrate complex provoking reduced ε-efficiency and/or a shift of the initial cut. In the same way, manipulating sites in TM-N might be communicated to the cleavage domain. The effect on ε-cleavage efficiency observed after glycine to alanine substitution at G33 (Higashide, Ishihara, Nobuhara, Ihara, & Funamoto, 2017) as well as the shifted cleavage pattern concomitant with the creation of an alternative ε-site at M51 in the G33Q mutant (Oestereich et al., 2015; Olsson et al., 2014) can be considered as direct hints to this communication. The propagation of effects arising from localized perturbations along the network of coupled protein fluctuations has been reported for several proteins even in the absence allosteric function (Bouguet-Bonnet & Buck, 2008; Clarkson et al., 2006; Ettayapuram Ramaprasad et al., 2017; Freire, 1999; Liu et al., 2006; Pan et al., 2000) and their general relation to disease has been discussed (Kucukkal, Petukh, Li, & Alexov, 2015). The synergy between binding and presentation of the ε-sites emerging from the current analysis is summarized in **Figure 12**. Surprisingly, the low local flexibility at the ε-sites remains intact, raising the question for an unfoldase activity of the enzyme. The rotation of the cleavage domain in the “bound-like” motion around a flexible joint upstream to the ε-site might even favour unwinding of the downstream P’-sites. Such a mechanism was suggested to contribute to initialization of hydrolysis (Bolduc et al., 2016).

**Figure 12.**
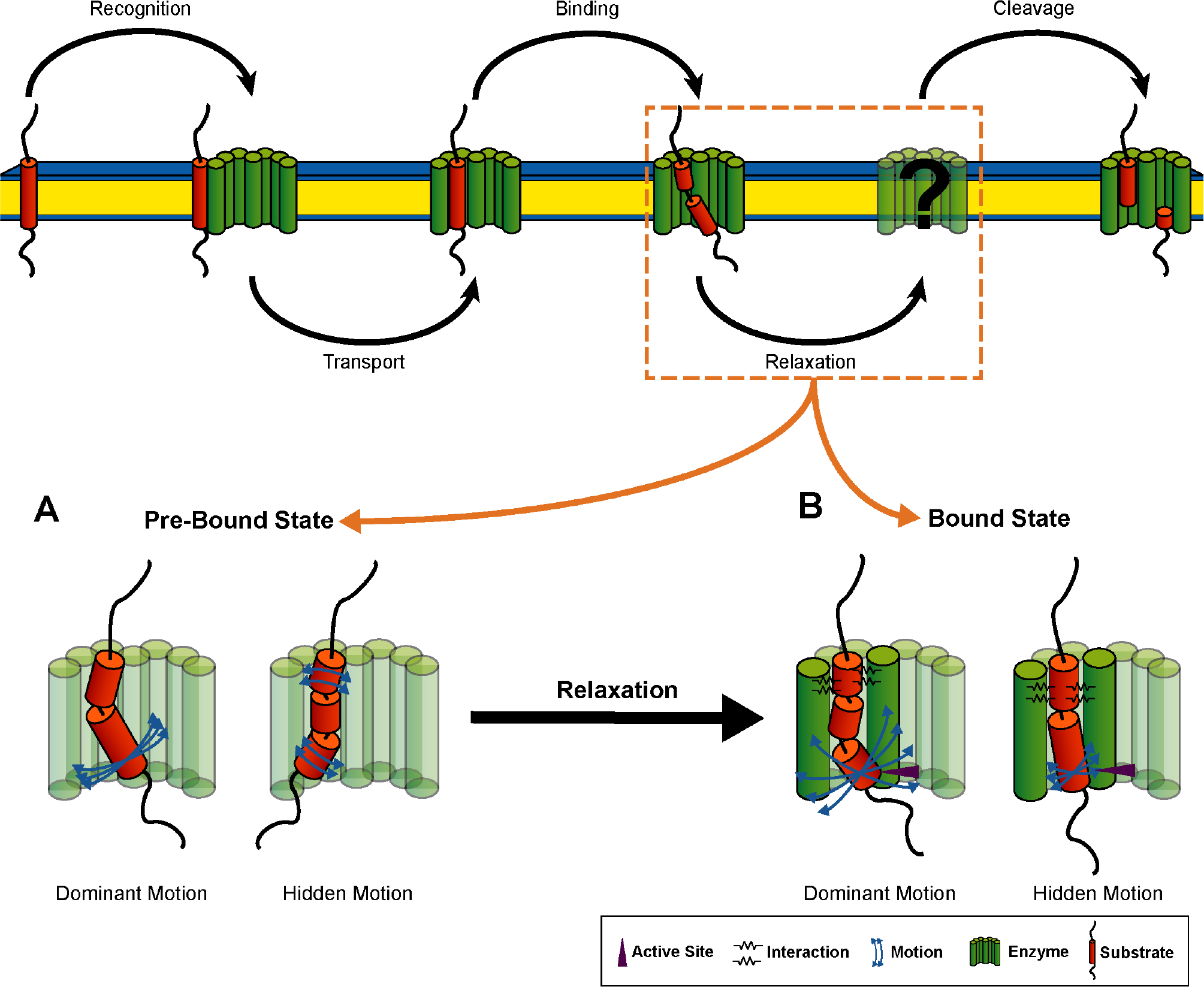
Binding-induced reorganization of C99 TMD conformational dynamics. (**A**) In the pre-bound state, large-scale bending coordinated by the G37G38 hinge prevails over low-amplitude motions where the helix flanks bend correlated with respect to the middle domain around hinge sites at V36G37 in TM-N and T43V44 in TM-C. Blue arrows indicate variability between backbone motions of WT and FADs. (**B**) Binding interactions (black springs) can selectively enhance this double-hinge motion, thus linking the region upstream to the ε-sites to catalytically important conformational relaxations. Consequently, softening or shifting the hinges in FAD mutants might affect the way in which the ε-cleavage sites interact with the enzyme’s active site (purple arrow).

While the repertoire of conformational motions of the substrate TMDs is mainly determined by their α-helical fold architecture, there is reason to suspect that the relative importance of individual motions reflects differences in sequence and local fold. The TMD of NOTCH, another prominent substrate of GSEC (Beel & Sanders, 2008), appears to be a straight helix (Deatherage et al., 2017) missing the large-scale bending flexibility that characterizes the C99 TMD. As a result, the TMDs of NOTCH and C99 adapt differently to the hydrophobic thickness of the membrane environment (Deatherage et al., 2017). In a similar manner, the coexistence of multiple dynamic features of substrate TMDs might determine how they adapt to interactions with the enzyme. Binding of different substrates might even take place at different interfaces, thus providing altered interactions (S. Li, Zhang, & Han, 2017). Therefore, relaxation after initial capture of the substrate might proceed along a series of intermediates of mutually adapted enzyme-substrate complexes (Ma & Nussinov, 2010). According to this model, neither dominance of a large-scale, collective motion nor local instability of an H-bond is, taken in isolation, a determining factors for GSEC cleavage, but rather their hierarchical organization (Henzler-Wildman et al., 2007). Generally, the corresponding transient steps are controlled by their kinetics in a gradually changing conformational landscape of the enzyme-substrate complex (Ma & Nussinov, 2010). The true binding-competent conformational changes might be identified only upon taking their kinetic differences into account. The GSEC cleavage of substrates with a different conformational dynamics signature of their TMDs suggests that the organization of the dynamic ensemble of the C99 TMD might not be a paradigm for the backbone dynamics of substrate TMDs in general. In our ongoing work, we compare the conformational dynamics of a large set of substrate TMDs in membrane environment as well as in the TFE/water mixture.

## Materials and Methods

### Peptide Sequences

Our model peptides consisted of residues 28-55 of C99 and carryied an additional KKW tag at the N-terminus (e.g. for wild-type: KKWK-GAIIGLMVGGVVIATVIVITLVML-KKK) as described previously (Pester, Barrett, et al., 2013; Pester, Götz, et al., 2013). The TMD of wild-type (WT) APP was compared to seven FAD mutants (T43I, V44A, V44M, I45T, I45F, V46I and V46F) which have previously been characterized with respect to cleavage efficiency, product-line preference and Aβ-ratios by several groups (Bolduc et al., 2016; Chávez-Gutiérrez et al., 2012; Chen et al., 2014; Dimitrov et al., 2013; Fernandez et al., 2016; Xu et al., 2016). The mutation site’s locations are depicted in Figure 1. Previously it was shown that similar C99 TMD-peptides are good substrates for GSEC (Chen et al., 2014; Yin et al., 2007).

### MD Simulations

Simulations were carried out as described in our previous work (Högel et al., 2018; Pester, Barrett, et al., 2013; Pester, Götz, et al., 2013; Scharnagl et al., 2014). Briefly, the peptides were solvated in a rectangular box (104 × 60 × 60 Å^3^), containing 80% 2,2,2-trifluoroethanol (TFE) and 20% water (v/v). After a 1.2 ns equilibration phase with gradual release of constraints on the peptides, 200 ns of free dynamics was conducted in a NPT ensemble (T = 293 K, p = 0.1 MPa). The last 150 ns of each trajectory were analyzed All simulations were performed using NAMD2.9 (Phillips et al., 2005) and the CHARMM22 force field with CMAP corrections (MacKerell et al., 1998).

### Analysis of structural and dynamic parameters

According to a geometrical criterion (Quint et al., 2010), a H-bond between the backbone amide hydrogen and the carbonyl oxygen is formed if the H⋯O distance is ≤ 2.6 Å and the N-H⋯O angle is within 180°±60°. We considered the H-bond emanating from the carbonyl oxygen at residue i to be closed if either the H-bond to NH_i+4_ (α-H-bond) or the H-bond to NH_i+3_ (3_10_ H-bond) was formed. The occupancy for each residue was computed as the fraction of closed H-bonds (α or 3_10_). For the calculation of water coordination numbers, water molecules with a hydrogen atom within 2.6 Å distance to the carbonyl oxygen atom of each residue were counted. This distance corresponds to the same cut-off as that used to define of H-bond occupancies. The packing score Si measures the contacts of the carbonyl oxygen of residue i to all other atoms j in the peptide by a cut-off-free approach (Grossfield, Feller, & Pitman, 2006). It is defined as the sum of the inverse of pairwise distances r_ij_ raised to the sixth power. Atoms belonging to the same residue i as the carbonyl were excluded. The rise per residue was calculated by a differential geometric approach (Guo et al., 2013) as used in our previous work (Högel et al., 2018; Scharnagl et al., 2014). The orientation of the cleavage domain was characterized by bending (θ) and swivel (Φ) angles of the helical segment carrying the cleavage sites (residues I47-M51) with respect to a helical segment located in TM-N (residues A30-L34, for definition of the angles see **Supplementary Figure 1**). Radial distribution functions and density distributions of water and TFE molecules were calculated using tools provided with the LOOS libraries (Romo, Leioatts, & Grossfield, 2014). Distances were measured between the centres of mass of side chains and solvent molecules. Mean structures were determined iteratively (Grossfield, Feller, & Pitman, 2007). Convergence of the simulations were investigated separately for a global and a local property (see **Supplementary Figure 2**). Backbone root mean-squared deviations (RMSD, Cα atoms of residues G29-M51) were calculated from mean structures in non-overlapping time windows with respect to the mean structure calculated over the full trajectory (Romo & Grossfield, 2011). The correlation times of H-bond fluctuations were calculated from the mean first passage of the autocorrelation functions through zero. The analysis indicates that local H-bond fluctuations as well as global backbone fluctuations for all simulations converge, at least after 30 ns. Therefore, standard errors of the mean for the presented properties were determined from block averaging using 5 non-overlapping blocks of 30 ns long blocks. Analysis was carried out using custom software based upon MDTraj (McGibbon et al., 2015). Graphical representations of structures were generated with VMD (Humphrey, Dalke, & Schulten, 1996).

### Dynamic domain analysis

The Dyndom program (Hayward & Lee, 2002) was used to analyse regions involved in hinge bending and twisting motions. Quasi-rigid domains of at least four residues were identified using a sliding window of 5 residues. The motions around flexible hinge sites were classified by the orientation of the screw axis relative to the helix axis. Motions with screw axis mainly perpendicular to the helix axis (%-closure >50%,) are classified as bending, while twisting motions have a screw axis mainly parallel to the helix axis (%-closure <50%) (Taylor et al., 2014). We subjected snapshots taken every 50 ps to analysis. The conformation with the lowest RMSD from the average structure was used as a common reference.

### Functional mode analysis

For functional mode analysis (FMA) we used the PLS-FMA program which uses a partial least-squares (PLS) model and was kindly provided by Bert de Groot (Krivobokova et al., 2012). Two functional order parameters were defined from the time series of the occupancies summed over H-bonds (α or 3_10_) in region I (emanating from carbonly-Os G33-G38, spanning residues G33-V42) and region II (emanating from carbonyl-Os V40-T43, spanning residues V40-I47), respectively. For the analysis we used all the heavy backbone atoms of residues K28-K54. The FMA model was constructed using 50% of the data and the remaining 50% were used for cross-validation. The required number of PLS components was determined from the dependence of the Pearson correlation coefficient between data and model (R_m_) as a function of the number of components. R_m_ converges to values >0.75 with 11-15 PLS components. For model building and cross-validation see **Supplementary Figure 3**. The structural changes that cause substantial variation in the order parameters were characterized by the ensemble-weighted, maximally correlated motions (ewMCM). For visualization, trajectories along the ewMCM vectors interpolating from low to high value of occupancies were used. To characterize the backbone motions in detail, two structures representing the conformational variability for low and high occupancies were extracted and analysed with DynDom (Hayward & Lee, 2002). To quantify similarities of ewMCM vectors from different peptides, the inner product of ewMCM vectors was calculated. The pairwise inner products of all peptides were used for hierarchical clustering (Kleiweg, 2014).

### Dynamics perturbation response analysis

Coupling between protein dynamics and binding can be well explained by an analytical model in which the reorganization of backbone dynamics is calculated as response to binding-induced interactions. Propagation of the perturbation throughout the protein is mediated by the correlation of residue fluctuations in the unperturbed state. Response of residues to perturbations upon binding has previously been formulated and analysed for several proteins and types of perturbations. In various studies the initial state’s average structure was perturbed by forces and the displacements of residues were monitored (Atilgan, Aykut, & Atilgan, 2011; Echave & Fernández, 2010; Ikeguchi, Ueno, Sato, & Kidera, 2005; Naritomi & Fuchigami, 2011). In other studies residue flexibility and connectivity were perturbed by adding interaction energies and the redistribution of conformational states was recorded (Erman, 2006; Ettayapuram Ramaprasad et al., 2017; Guarnera & Berezovsky, 2016; Ming & Wall, 2006; Zheng & Brooks, 2005; Zheng et al., 2007). Here we applied dynamic perturbations, because (i) the mean structures of the TMDs did not show substantial conformational changes even for the mutants, (ii) a reasonable model for functionally relevant forces was missing, and (iii) given the importance of the direction of perturbing forces (Atilgan et al., 2011) applying random kicks might not by sufficiently informative enough. In perturbation-response calculations, the dynamics of the unperturbed protein is often derived from an elastic network description. Here, we formulated a quasi-harmonic extension of this concept including non-harmonicities in the variance-covariance matrix from MD, as outlined below. The enzyme as well as its interactions with the substrate TMD were treated implicitly using concepts from Gaussian Network Models (Erman, 2006; Guarnera & Berezovsky, 2016). The enzyme was assumed to constitute a linear chain of binding sites. Linear connectivity was accounted for by strong attractive spring constants κ between nearest neighbours. The non-covalent interactions between substrate TMD and enzyme were treated as a source of elastic energy that influences the motions of the explicitly treated mechanical degrees of freedom which are the Cα atoms of the helical core of the TMD. Weak attractive spring constants γ harmonically and isotropically restrain the binding sites. The binding-induced reorganization of conformational dynamics was characterized by the variance-covariance matrix of the perturbed TMD calculated using techniques for partitioned, interacting subsystems (Ming & Wall, 2005a; Woodcock et al., 2008).

We started by setting up the potential energy of the interacting TMD-enzyme system in the form of a combined Hessian matrix in block form, as exemplified in Equation 1 for the interaction of one binding site on the TMD (Cα atom of residue i) with binding site m on the enzyme. We assumed that each residue of the TMD interacts with only one binding site of the enzyme and exclude multiple binding to the same site:

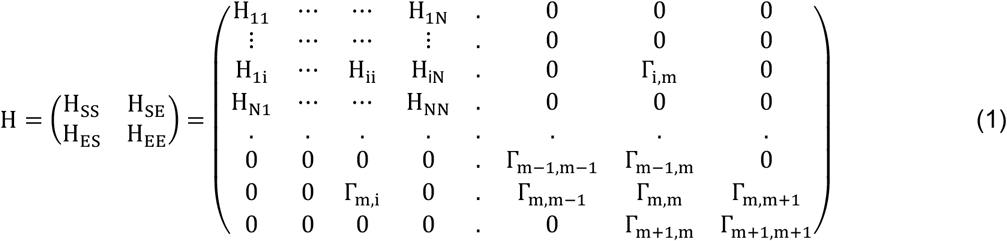

The Hessian of the free substrate TMD (H^u^_ss_) is a NxN matrix of 3×3 sub-matrices (N = number of Cα atoms). Formally, H^u^_ss_ is written as pseudo-inverse of the quasi-harmonic variance-covariance matrix C^u^_ss_ of the unperturbed system (k_B_ = Boltzmann constant, T = temperature): 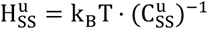. Due to the interactions with the enzyme, the diagonal super-elements H_ii_ of H_ss_ differ from the unperturbed ones. These can be calculated from the condition, that the sum of elements of each row and column in the interacting system must be zero, i.e. 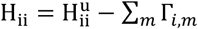 and 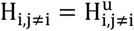. The harmonic and isotropic interaction between atom i on the TMD and enzyme site m is characterized by the spring constant γ and off-diagonal 3×3 sub-matrices Γ_*i,m*_ entering the interaction matrix H_SE_ (H_ES_ is the transposed of H_SE_):

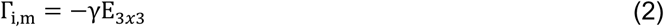

The Hessian H_EE_ describing the enzyme with M binding sites is a MxM tri-diagonal matrix of 3×3 sub-matrices. Each enzyme site m interacts with its nearest neighbours m±1 in the linear chain. Interaction strength is given be the spring constant *κ* and interactions are assumed to be isotropic. Therefore, H_EE_ contains off-diagonal submatrices:

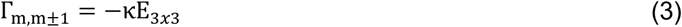

The diagonal submatrices Γ_*m,m*_ of H_EE_ take all interactions into account:

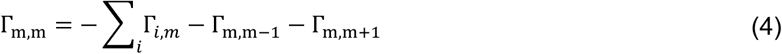

By integrating out the environment (Ming & Wall, 2005a; Woodcock et al., 2008), the effective Hessian H^b^ss for the peptide under the influence of binding-induced dynamic perturbations can be calculated as:

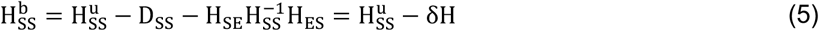

Thereby, D_ss_ is a 3Nx3N diagonal matrix accounting for the differences between H_ii_ and H^u^ii. Finally, the covariance matrix of the perturbed peptide is recovered from inversion of H^b^ss:

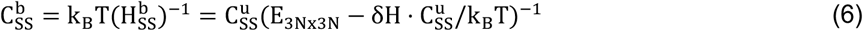

where E_3Nx3N_ is a 3Nx3N unit matrix. Here, δH accounts for the harmonic interaction with the enzyme. The interaction is propagated in the peptide along the correlated fluctuations in the unperturbed state, described by C^u^_ss_, utilizing the full dynamic information from MD. As a result of substrate-enzyme binding, the peptides have changed MsFs (diagonal elements of C^b^_ss_) and a reorganized space of backbone motions, determined from an eigenvector decomposition of C^b^_ss_ and C^u^_ss_, respectively. The essential sub-space was defined to describe 85% of the overall Cα mean-square fluctuations. The overlap between the functional modes and the essential sub-space, as computed by Equation 7, quantifies similarity of the functional mode (FM) and the essential sub-spaces (*ν*_*i*_) of the free and bound states, respectively. The number of modes necessary to describe 85% of Cα MSFs (n) varies between 6 and 10. For perturbation-response mapping custom-built software was developed in Fortran90.

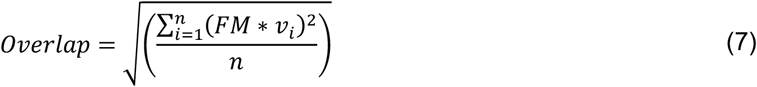

The value of the spring constant *γ* was determined from the condition that in the linear response limit the change in MSF is proportional (i) to the square of the MSF in the unperturbed state and (ii) to the perturbation, i.e. the spring constant (K. sato, Ito, Yomo, & Kaneko, 2003). We therefore calculated the relative change of MSF, ΔMSF, for spring constants *γ* between 1 and 10 kcal/(mol Å^2^).

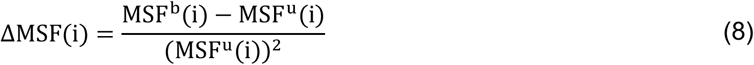

In the linear regime, the slope ΔMSF/Δγ should be constant. We tested this relationship for several residues as well as for the overall MSF in the binding model with the strongest interactions (see **Supplementary Figure 4)**. We concluded that a spring constant γ = 5 kcal/(mol.Å^2^) provides a good approximation to the linear regime. The value is in the lower limit of weak intra-helical i,i+4 force constants (Moritsugu & Smith, 2007; Pester, Barrett, et al., 2013). A much stronger spring constant describes the “covalent” bonds between enzyme sites. Here we use *κ* = 150 kcal/(mol Å^2^).

### Normalized fluctuation profiles and Z_score_ analysis

Changes in MSF between two models were quantified by first calculating the normalized fluctuation profile for each model and, second, taking the Z-score of the difference of the normalized profiles as described previously (Gu & Bourne, 2007). For each interaction model, the normalized fluctuation profile F_norm_ was determined excluding outliers (M_score_ > 3.5) when calculating the mean μ_MSF_ and standard deviation σ_MSF_ of peptide fluctuations:

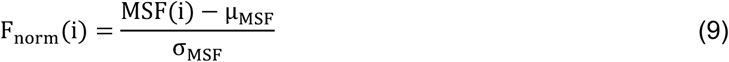

The difference ΔF between normalized fluctuations was calculated in order to identify residues with significant changes in fluctuations. For this difference, the score Z_diff_ was calculated:

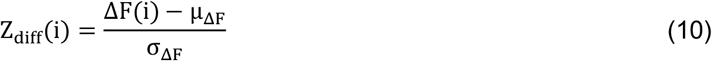

Values >0 (<0) indicate above (below) average fluctuation change. Changes were regarded as significant if they differed by more than one standard deviation from the mean.

## Supporting Information

Supporting information contains Supplementary Figures 1 - 5, 8.1 and 9.1

## Acknowledgements

The authors would like to thank Dieter Langosch for stimulating discussions and Norman Blümel for assistance with simulation setup and production runs. This work was supported by the Deutsche Forschungsgemeinschaft (DFG) within the framework of the DFG research unit FOR2290 (Grant SCHA630/4-1). Computing resources were provided by the Leibniz supercomputing Centre (LRZ) through grants ta511 (Linux Cluster) and pr42ri (SuperMUC).

## Author Contributions

AG and CS designed research, AG performed research, CS contributed new analysis tools. Both authors analysed data, wrote the manuscript and have given approval to its final version.

